# Non-coding rare variant associations with blood traits on 166 740 UK Biobank genomes

**DOI:** 10.1101/2023.12.01.569422

**Authors:** Diogo M. Ribeiro, Olivier Delaneau

## Abstract

Large biobanks with whole-genome sequencing now enable the association of non-coding rare variants with complex human traits. Given that >98% of the genome is available for exploration, the selection of non-coding variants remains a critical yet unresolved challenge in these analyses. Here, we leverage knowledge of blood gene regulation and deleteriousness scores to select non-coding variants pertinent for association with blood-related traits. We leverage whole genome sequencing and 59 blood cell count and biomarker measurements for 166 740 UK Biobank samples to perform variant collapsing tests. We identified hundreds of gene-trait associations involving non-coding variants across the 59 traits. However, we demonstrate that the majority of these non-coding rare variant associations (i) reproduce associations known from common variant studies and (ii) are driven by linkage disequilibrium between nearby common and rare variants. This study underscores the prevailing challenges in rare variant analysis and the need for caution when interpreting non-coding rare variant association results.

## Introduction

Despite the large success of genome-wide association studies (GWAS) in linking variants to traits, these primarily rely on common variants assessed in SNP arrays, thus often identifying variants in linkage disequilibrium (LD) with causal variants^1^. While genotype imputation from fully sequenced reference panels expands the variant repertoire in GWAS studies^2,3^, SNP-based heritability estimates for traits still fall short of pedigree-based heritability^4^. A possible reason for this is that the causal variants are rare variants (e.g. minor allele frequency, MAF < 1%) not included in GWAS studies. Consequently, rare variant analyses are thought to provide biological insights complementary to those of GWAS and account for part of the missing heritability^5,6^. As natural selection shapes the distribution of effect size and allele frequency of variants associated with complex traits, rare variants tend to confer larger effect sizes compared to common variants^7^. In other words, alleles that strongly predispose individuals to disease are likely to be highly deleterious and are thus kept at low frequencies in the population through purifying selection. The study of rare variation holds great promise, enabling clinical studies in individual patients and is poised to reveal novel gene targets for pharmaceutical intervention^8–10^. For instance, the discovery of rare variants in the PCSK9 gene in individuals displaying low LDL cholesterol levels led to clinical trials and the subsequent approval of a PCSK9 inhibitor to treat cardiovascular disease^11,12^.

Current rare variation studies have analysed hundreds of thousands of individuals and linked hundreds of variants and genes to human traits and diseases^8–10,13^. However, most rare variants studies so far have focused on the coding regions of the genome, primarily owing to the cost-effectiveness and widespread availability of whole exome-sequencing (WES) compared to whole-genome sequencing (WGS). Moreover, coding variants have a straightforward interpretation (e.g. altering protein sequence) and potential high effect on the phenotype. Nevertheless, exonic regions represent only ∼2% of the human genome, whereas up to 88% variants associated with complex traits and diseases through GWAS were found in the non-coding regulatory regions of the genome^14,15^. Yet, large-scale studies associating non-coding rare variation with phenotypes are scarce^16,17^. This is largely due to the requirement of WGS data in large cohorts and the challenging interpretation of non-coding variant effects^18^. Several biobanks have recently made efforts to include WGS data, as is the case of the UK biobank^19,20^, TopMed^21^, and the All of Us^22^ biobanks. These datasets allow studying the effect of hundreds of millions of variants towards thousands of available phenotypes. Thus, the remaining challenge now relates to developing approaches that efficiently explore the extensive search-space of the non-coding portion of the human genome.

The spatio-temporal and tissue-specific expression of genes is achieved by nearby non-coding regulatory elements – usually within 1Mb of the transcription start site (TSS) – including enhancers, promoters and insulators^23^. Variants in these regulatory regions are expected to influence traits and diseases by modulating gene expression levels^24,25^. While promising, defining gene regulatory regions is challenging due to the incomplete knowledge of the regulatory architecture for each gene^26^ and gene regulation being highly cell-type and context-specific^27^. Despite this, substantial efforts have been invested in the non-trivial task of identifying regulatory regions and their target genes, including through (i) chromatin contacts from Hi-C data^28,29^, (ii) correlating gene expression with enhancer activity (e.g. H3K27ac and H3K4me1 histone modifications) across individuals^30^ and various cell types^31,32^ and (iii) employing integrated models that combine multiomics data^33,34^. Recent studies have harnessed this information, along with expression quantitative trait loci (eQTL) and single cell ATAC-seq data to link non-coding GWAS variants to genes^35,36^. Crucially, these gene-based approaches allow the identification of genes mediating the link from non-coding variation to phenotype – a key limitation in GWAS studies. Translating such approaches to the context of rare variants holds great promise and represents a compelling research avenue.

Here, we leverage 166 740 UK Biobank whole genome sequences to perform a genome-wide association analysis between non-coding rare variants and 59 continuous blood traits. We explore several orthogonal gene regulatory annotations in guiding variant region choice, (i) cis-regulatory domains for four blood cell types derived from ChIP-seq and RNA-seq data across hundreds of individuals^30,37^, (ii) activity-by-contact (ABC) model gene-enhancer links across 60 blood cell types derived from multiomics data^33,34^ and (iii) promoter-capture Hi-C data across 17 blood cell types^29^. Across these regulatory annotations, we identified 1406 gene-trait associations, demonstrating a comparable discovery power to coding regions. However, we show that the majority of these rare variant associations are not independent from known common variants, with many associated rare variants in linkage disequilibrium with GWAS variants. Our results highlight the need for caution when interpreting non-coding rare variant associations.

## Results

### Regulatory annotations linking non-coding variants to genes

Our approach to exploring associations between non-coding rare variants and traits takes a gene-centric perspective. Given the lack of consensus approaches to identify regulatory regions and link them to genes, we evaluate the utility of three distinct, complementary approaches to map rare non-coding variants to protein-coding genes (**Figure 1**). First, we gather cis-regulatory domains (CRDs) linking 118 408 regulatory domains and 8 991 protein-coding genes (**Table 1**). These CRDs are inferred from ChIP-seq data, particularly for histone modifications indicative of active enhancers and promoters (H3K4me1 and H3K27ac), as well as RNA-seq data obtained from a cohort of individuals, focusing on four key blood cell types (monocytes, neutrophils, T cells, and lymphoblastoid cell lines)^30,37^. Second, we obtained 601 733 gene-enhancer links for 13 516 genes from the activity-by-contact (ABC) model, which leverages Hi-C, ChIP–seq and ATAC-seq data, for a set of 60 blood cell types (ABC score > 0.1)^33,34^. Third, we exploit 808 237 gene-chromatin links to 14 638 genes, from promoter capture Hi-C (PCHi-C, hereafter referred to as HIC) data produced on 17 primary blood cell types by Javierre et al. 2016^29^, only considering links within 1Mb of the gene transcription start site with CHiCAGO score > 5 (see Methods).

**Figure 1.**
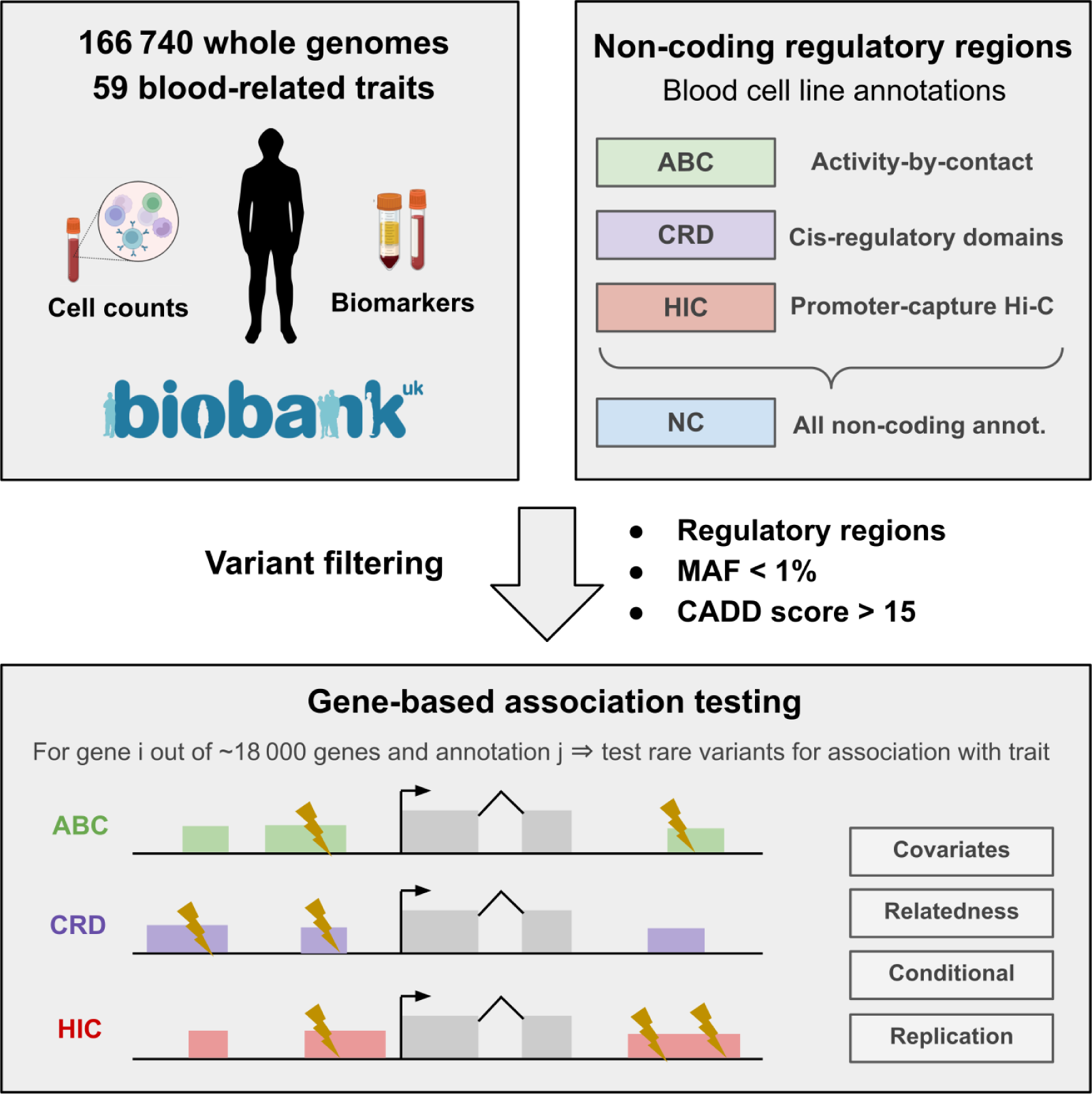
Schematic representation of the study approach. We outline our comprehensive approach to investigating non-coding genetic variants and their associations with 59 blood-related phenotypes in a cohort of white British individuals from the UK Biobank. To explore non-coding regions for potential trait associations, our focus is on gene regulatory regions derived from three distinct datasets: ABC (Activity-by-contact), CRD (cis-regulatory domains) and HIC (promoter capture Hi-C, PCHiC), spanning various blood cell types. In certain analysis, these three annotations are combined and referred to as the NC (non-coding) annotation. Genetic variants were filtered for MAF < 1% and CADD (Combined Annotation Dependent Depletion) score > 15. REGENIE was used for gene-based association tests (SKAT-O tests), including covariates and relatedness estimates. These were performed per gene per trait and per annotation (e.g. ∼18 000 genes × 59 traits × 3 annotations). Additionally, follow-up analyses encompassed replication assessments and conditional analyses of GWAS common variants.

**Table 1.** Summary of association tests performed across annotations on 59 UK Biobank blood-related traits. FDR is based on the Benjamini-Hochberg procedure from SKAT-O association p-values. Conditional results are on a subset of 42 traits. Only associations significant before conditioning and remaining significant after conditioning at the same threshold are counted.

| Annotation | Genes tested | Median SNPs | Gene-trait FDR < 5% Before conditional | Gene-trait P < 1e<sup>-9</sup> Before conditional | Gene-trait FDR < 5% Remain after conditional | Gene-trait P < 1e<sup>-9</sup> Remain after conditional |
| --- | --- | --- | --- | --- | --- | --- |
| CRD | 8991 | 243 | 391 | 128 | — | — |
| ABC | 13 983 | 80 | 142 | 27 | — | — |
| HIC | 14 638 | 462 | 1026 | 320 | — | — |
| NC | 16 673 | 580 | 1406 | 380 | 88 | 9 |
| CDS | 18 838 | 165 | 964 | 279 | 276 | 76 |

Since most genetic variants are expected to have no discernible impact on phenotypes, rare variant association methods benefit from a pre-selection of pertinent variants to include in statistical testing^18,38^. For this, we used Combined Annotation Dependent Depletion (CADD) scores^39^, which leverage metrics of non-coding sequence functionality such as sequence conservation, chromatin state and molecular quantitative trait loci to predict pathogenicity for each SNP. We thus considered only rare SNPs (MAF < 1%) with CADD score > 15, which is equivalent to retaining the ∼3.1% most deleterious variants in the genome. After this filter, we retained 2 191 590, 1 062 230 and 11 155 962 autosomal SNPs for the CRD, ABC and HIC annotations, respectively. The variation in SNP numbers across annotations is inherent to the diverse data types, models, and cell types used. Additionally, the number of SNPs tested per gene is also highly variable, with a median of 243 SNPs in the CRD annotation, 80 for ABC model and 462 for HIC annotations (**Table 1**, **Supplementary Figure 1**). By comparison, the coding sequence (CDS) of genes feature a median of 165 variants. It is essential to note that regulatory variants overlapping with the CDS regions of any gene were deliberately excluded in non-coding rare variant analysis (see Methods).

### Non-coding rare variant association to traits

We performed gene-centric SKAT-O tests for each of the three regulatory annotations separately using REGENIE^40^, which allows us to account for relatedness, population structure and other covariates (e.g. sex, age). We considered quality-controlled genotypes from the whole genome sequencing data from the UK Biobank, across 166 740 white British individuals (see Methods). To understand what can be obtained through rare variant associations, we first present association results without conditioning for common variants. In later sections we present results conditioning for known GWAS variants for a subset of traits with available data. Therefore, we started by performing unadjusted association tests against 59 continuous traits, comprising all 29 blood cell counts (e.g. monocyte count) and 30 blood biomarkers (e.g. vitamin D, bilirubin) available in the UK Biobank (**Supplementary Table 1**). Blood-related phenotypes were deliberately chosen to be aligned with the blood regulatory annotations available, in order to tackle the issue of tissue and cell-type specificity of gene regulation.

Because we conduct only one test per gene (ranging from 8 991 to 14 638 genes per annotation), gene-based association tests exhibit a reduced multiple testing burden compared to single variant tests, such as those in GWAS. We therefore considered significant associations those having an FDR < 5%. This FDR threshold corresponds to SKAT-O p-values ranging from 8.7e^-6^ to 3.7e^-5^, depending on the specific annotation. Across all traits tested, we found between 142 and 1026 significant gene-trait associations — involving 90 to 510 genes and 35 to 56 traits, across the three different regulatory annotations tested (**Table 1**, **Figure 2**, FDR < 5%). The HIC annotation (highest number of variants) provided the most discoveries, while the ABC annotation (lowest number of variants) yielded the fewest. Imposing a more stringent p-value cutoff (P<1e^-9^) we find between 27 and 320 gene trait associations (18 to 151 genes, 15 to 44 traits, **Supplementary Table 2**). Overall, the three non-coding annotations used show to be highly orthogonal, with only 18 (1.3%) gene-trait associations replicated across the three annotations and between 31 and 57 associations being consistently replicated across two annotations (**Supplementary Figure 2**).

**Figure 2.**
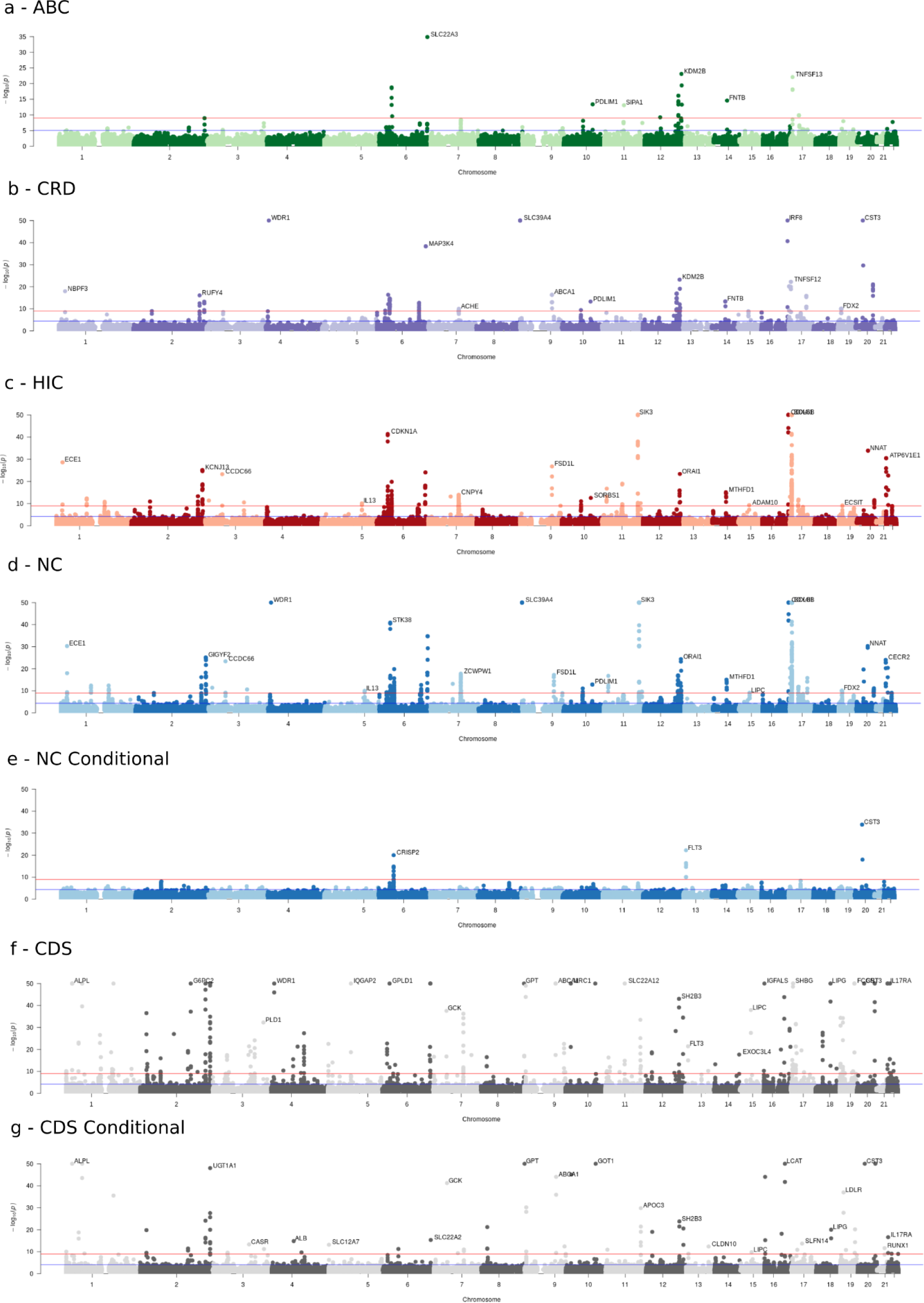
Manhattan plot for gene-based rare variant associations across 59 traits and multiple annotations. (a) ABC, (b) CRD, (c) HIC, (d) and (e) NC, (f) and (g) CDS. Each dot represents a gene-trait pair across chromosome positions (x-axis, gene TSS position used), with -log_10_ p-values (capped at 50) of the SKAT-O tests in the y-axis. Subplots (e) and (g) represent association results when conditioning on GWAS common variants across 42 traits (see below). Red lines represent P<1e^-9^ cutoff, blue lines represent the FDR<5% cutoff (calculated for each annotation separately). Gene symbol labels for the most strongly associated gene per chromosome were added.

To contextualise these findings within the scope of CDS regions utilising the same approach (e.g. SNPs with CADD > 15), we first combined SNPs from the three regulatory annotations into a single non-coding (‘NC’) mask. We found that non-coding regions discover a larger number of associations than CDS regions, with 1406 gene-trait associations from non-coding regions and 964 associations from coding regions (FDR < 5%, **Table 1**, **Figure 2**). Non-coding regions also identify a higher number of high-confidence associations (P<1e^-9^) than CDS regions (380 vs 279 gene-trait associations). However, when inspecting gene-trait associations between NC and CDS we notice a higher number of “gene-trait peaks” for NC compared to CDS (**Figure 2**). This phenomenon is attributed to the intricate nature of non-coding regulatory elements, which often have the capacity to influence multiple genes^41^. Consequently, the same non-coding SNPs are tested for association with multiple genes, thus leading to the association of multiple genes from a single signal. Such overlap does not occur in CDS regions, where typically SNPs relate to only one coding gene. For instance, out of 707 genes associated with traits in the NC annotation (FDR < 5%), only 242 of them (34.2%) had no other associated gene within a 1Mb radius. In contrast, this was true for 314 out of 481 (65.2%) of the CDS-associated genes. A similar pattern persisted for gene-trait associations with P<1e^-9^, where only 64 out of 187 (34.2%) NC gene-trait associations exhibited no other nearby associations, whereas this is true for 103 out of 150 (68.8%) for CDS gene-trait associations.

Although we found non-coding associations for 57 of the 59 traits tested (e.g. NC annotation FDR 5%), some traits displayed a much higher number of associations (e.g. 132 genes associated with mean platelet volume) than others (e.g. 1 gene associated with glucose levels, **Supplementary Figure 3**). Indeed, traits had a median of 20 gene associations through non-coding rare variants, compared to 17 through CDS variants (**Supplementary Figure 4**). The number of genes associated with traits through non-coding variants is significantly correlated with the number of genes known to be associated with those traits in the GWAS catalog (Pearson correlation = 0.37, p-value = 5.2e^-3^) and Genebass datasets (Pearson correlation = 0.69, p-value = 2.1e^-9^). This suggests that our findings align with existing knowledge and expected trait heritability.

In order to assess the effectiveness of utilising non-coding annotations based on experimental datasets, we generated a ‘control’ annotation. To construct this control set, we initially determined the number of variants associated with each gene within the ‘NC’ annotation. Subsequently, we randomly selected an equivalent number of variants for each gene from a pool of variants adhering to specific criteria: (i) residing within a 1Mb radius of each gene’s transcription start site (TSS), (ii) possessing a CADD score exceeding 15, and (iii) not being part of either the ‘NC’ or ‘CDS’ annotation (see Methods). In this manner, we obtained a control set of variants for ∼95% of the genes tested in ‘NC’. We then compared the discovery power of variants from 15 687 genes between ‘NC’ and the matching control variant set across the 59 traits. The control set identified 340 and 101 gene-trait associations at FDR 5% and P<1e^-9^, respectively (**Supplementary Table 3**). This represents a 2.8 to 3.2-fold decrease in discovery power compared to ‘NC’ variants, underscoring the significance of employing biologically-relevant regulatory annotations in enhancing the discovery power. However, finding hundreds of significant gene-trait associations with randomly sampled variants also suggests that the functional annotations used are incomplete, or that other cell-types and tissues could be pertinent to the assessed traits. Moreover, control variants may be coincidentally linked to a causal gene in the region, since the genomic windows used to sample variants may sometimes encompass dozens of genes.

### Out-of-sample replication and sample size

Next, we evaluated the replication rate of gene-trait associations across different samples. For this, we randomly split the set of 166 740 WGS samples into two distinct groups: a discovery set of 136 740 samples and a replication set of 26 212 samples, excluding related samples between the two groups (see Methods). First, we note that the replication set of 26 212 samples lacks the power required to uncover gene-trait associations with rare variants, as evident from the discovery of only 50 NC associations at FDR < 5% (72 for CDS annotation). We then used the two sets of samples in a discovery/replication approach. We found that 41.3% of 1113 NC gene-trait associations identified with the 136 740 samples were replicated at a significance level of P < 0.05 in the 26 212 samples subset, whereas only 4.3% were expected to replicate by chance (**Figure 3a**). This percentage increased to 70.7% when considering gene-trait associations with P<1e^-9^ in the discovery dataset, an expected increase given these comprise stronger associations. In comparison, CDS regions exhibited a higher degree of associations that could be replicated, with replication rates ranging from 47.8% (FDR<5%) to 86.5% (P<1e^-9^). Moreover, association p-values across all tested gene-trait pairs (>980 000 gene-trait pairs) were moderately correlated between the 136 740 and 26 212 sample sets (**Supplementary Figure 5**, Spearman correlation = 0.39 for NC and 0.41 for CDS, p-value < 2.2e^-16^). These results underscore the consistency and reliability of the associations derived from this analysis.

**Figure 3.**
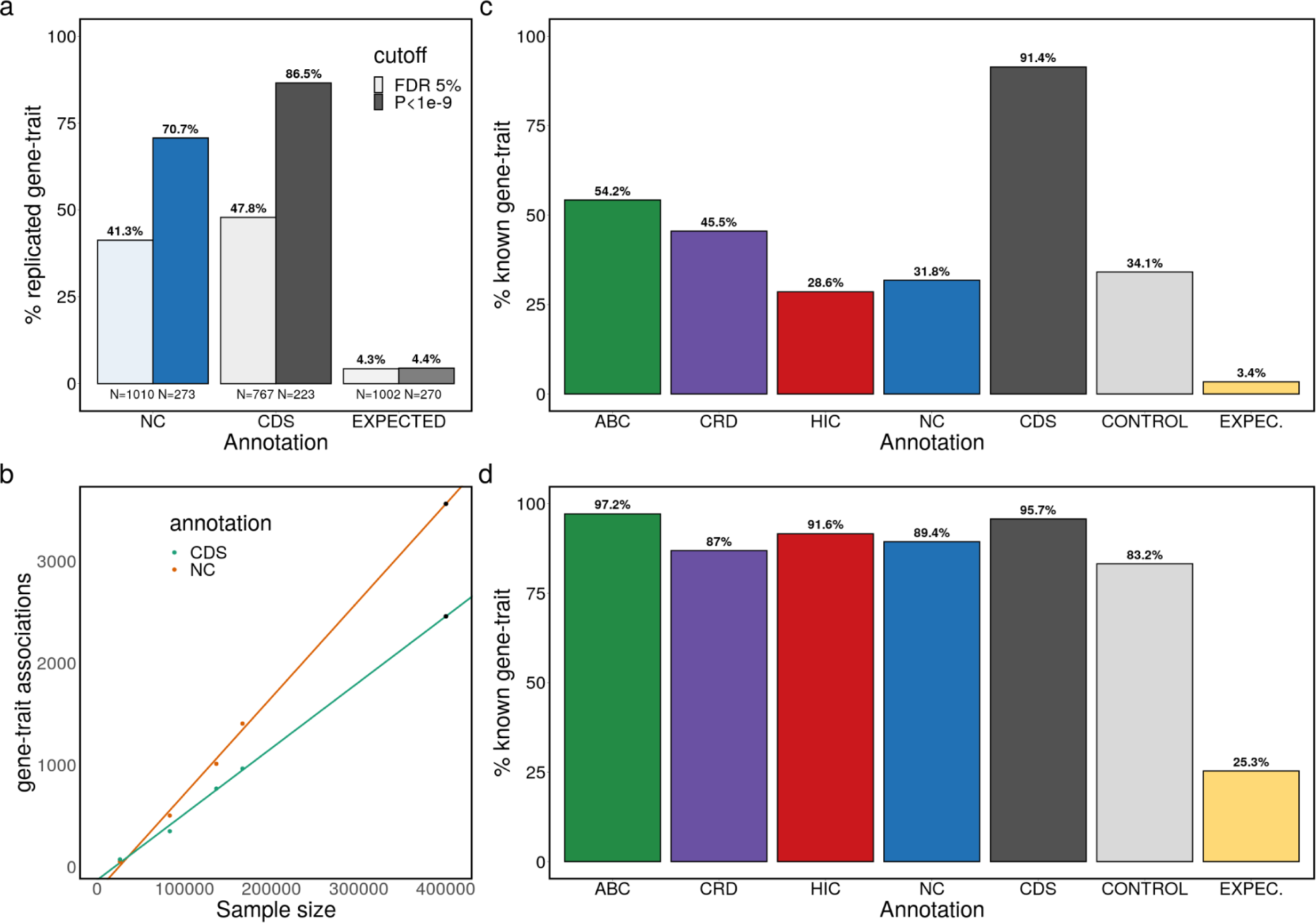
Replication of discovered gene-trait associations. (a) percentage of gene-trait pairs discovered in a set of 136 740 samples that have an association p-value < 0.05 in 26 212 samples, depending on annotation and discovery p-value cutoff; (b) correspondence between sample size and number of discovered gene-trait associations at FDR 5%. The black dot at 400 000 samples is a prediction based on the previous four samples sizes tested; (c) percentage of gene-trait pairs at FDR 5% significance present on GWAS catalog or Genebass datasets depending on annotation; (d) percentage of gene-trait pairs at FDR 5% present on GWAS catalog or Genebass either through direct gene-trait associations or indirect associations within a 1Mb window. ‘EXPEC.’ refers to a random shuffling of NC annotation association p-values.

Given the large difference in significant associations depending on sample size, we examined the relationship between sample size and number of associations discovered. Assessing discovery when downsampling to 26 212, 83 370, 136 740 and 166 740 samples, we found a strong linear relationship between these (**Figure 3b**, NC correlation = 0.99, p-value = 4.8e^-3^; CDS correlation = 0.99, p-value = 5.2e^-3^). Focusing on the sets of 166 740 and 83 370 samples, which represent a 2-fold sample size difference, we discovered 2.8 times more gene-trait associations with the larger sample size (both for NC and CDS). Moreover, these results indicate that discovery is far from plateauing, with an expected 3565 NC gene-trait associations discovered at FDR 5% (2460 for CDS) with 400 000 samples, which corresponds to the full number of British samples with WGS data in the UK Biobank. A similar relationship is expected for associations with P<1e^-9^, with 933 NC gene-trait associations expected for 400 000 samples (**Supplementary Figure 6**).

### Most rare variant associations to traits are known

We assessed whether the non-coding gene-trait associations discovered here matched discoveries from common variants (GWAS) and coding rare variants (exome-seq). For instance, we identified a non-coding rare variant association between CST3 (Cystatin C) gene and blood Cystatin C levels (SKAT-O p-value = 1.7e^-75^ for CRD annotation). Moreover, we found associations between the IRF8 gene and monocyte counts (SKAT-O p-value < 5.7e^-61^ for CRD and HIC annotations), a well documented association^42,43^. To comprehensively evaluate our ability to identify known gene-trait associations, for each of the 59 traits assessed, we gathered gene-trait associations from two datasets: (i) GWAS catalog, which is largely composed on common variant analyses, and (ii) Genebass database, which used coding rare variants from UK Biobank exome-seq data (see Methods). We found that between 28.6% (HIC) and 54.2% (ABC) gene-trait pairs at FDR<5% were previously documented in either the GWAS catalog or Genebass databases, depending on the non-coding annotation used (**Figure 3c**). This percentage increased to between 35.9% (HIC) and 92.6% (ABC) when considering gene-traits associations with P<1e^-9^ (**Supplementary Figure 7**). This overlap with known gene-trait pairs far exceeds the 3.4% to 3.7% expected by chance, as determined by shuffling SKAT-O p-values for gene-trait associations based on the NC annotation. The vast majority of CDS gene-trait associations (>91.4%) are known, however, this is expected as the Genebass database uses rare coding variants from the UK Biobank. Indeed, when considering only gene-trait pairs from the GWAS catalog, the replication rate drops to 67.2% (compared to 88.7% in Genebass, FDR 5%, **Supplementary Figure 8**). Intriguingly, using the control set of non-coding variants (same number of variants as NC) produces similar rates of replication as NC (34.1% with FDR < 5%, 43.6.% with P<1e^-9^). This suggests that the control variants may reveal *bona fide* gene-trait relationships with a false discovery rate similar to the regulatory regions – just with a lower number of discoveries, as seen previously.

Given that non-coding variants have the capacity to influence several nearby genes through mechanisms such as enhancers targeting multiple genes, we extended our analysis to encompass nearby genes. Instead of merely counting a match when the exact gene-trait pair was previously documented, we broadened our search to include cases where another gene within a 1Mb window matched the known gene-trait pair (see Methods). Doing so, we found that 87% (CRD) to 97.2% (ABC) of gene-trait pairs match, similar to what is observed for CDS annotations (95.7%, **Figure 3d, Supplementary Figure 7**). While this expanded scan is expected to increase the proportion of matches found, we note that the expected overlap value when applying random shuffling is 25.3%. These findings underscore that, while non-coding variant analysis presents challenges in pinpointing the exact causal gene compared to coding variants, it adeptly identifies likely causal regions, providing valuable insights into the genetic underpinnings of traits.

### Most rare variant associations are not independent of common variants

Given that most gene-trait associations identified here are known from previous studies, we next addressed whether these gene-trait associations are independent from known common variants. For this, we used GWAS summary statistics from the Pan-UK Biobank study across 42 of the 59 traits assessed (see Methods, **Supplementary Table 1**). We derived independent GWAS hits at P<1e^-7^ using the GCTA-cojo joint model^44^. Subsequently, we conducted REGENIE gene-based association tests conditioning on all independent GWAS hits per trait (median of 968.5 variants, ranging from 201 to 2164). We found that the vast majority of gene-trait associations were not independent of common variants, as only 88 out of 929 (9.47%) NC gene-trait associations at FDR<5% passed this significance level after conditioning for common variants (**Figure 4a**, **Figure 2e**, **Table 1**). Furthermore, when focusing on the associations with P<1e^-9^, only 9 out of 242 (3.72%) gene-trait associations were preserved (**Supplementary Table 4**). These 9 associations include well known associations, such as the Cystatin C gene (CST3) association with Cystatin C levels (SKAT-O p-value before conditional = 2.1e^-20^, after conditional = 1.3e^-30^) and several cysteine-rich secretory protein genes (CRISP1, CRISP2 and CRISP3) associated with ‘Mean sphered cell volume’ and other reticulocyte phenotypes. These three CRISP genes are clustered together in chromosome 6 and at least CRISP2 is reported to be associated with reticulocyte phenotypes^45^. Although most previous associations drop in significance, conditional analysis also reveals several associations that were not significant before conditioning. We therefore report 316 gene-trait associations at FDR<5% independent of common variants, across the 42 traits (i.e. 228 additional associations compared to the 88 reported above). However, only 15 of these associations (i.e. 6 additional, compared to the 9 reported above) pass the P<1e^-9^ threshold (**Figure 4a, Supplementary Table 4**).

**Figure 4.**
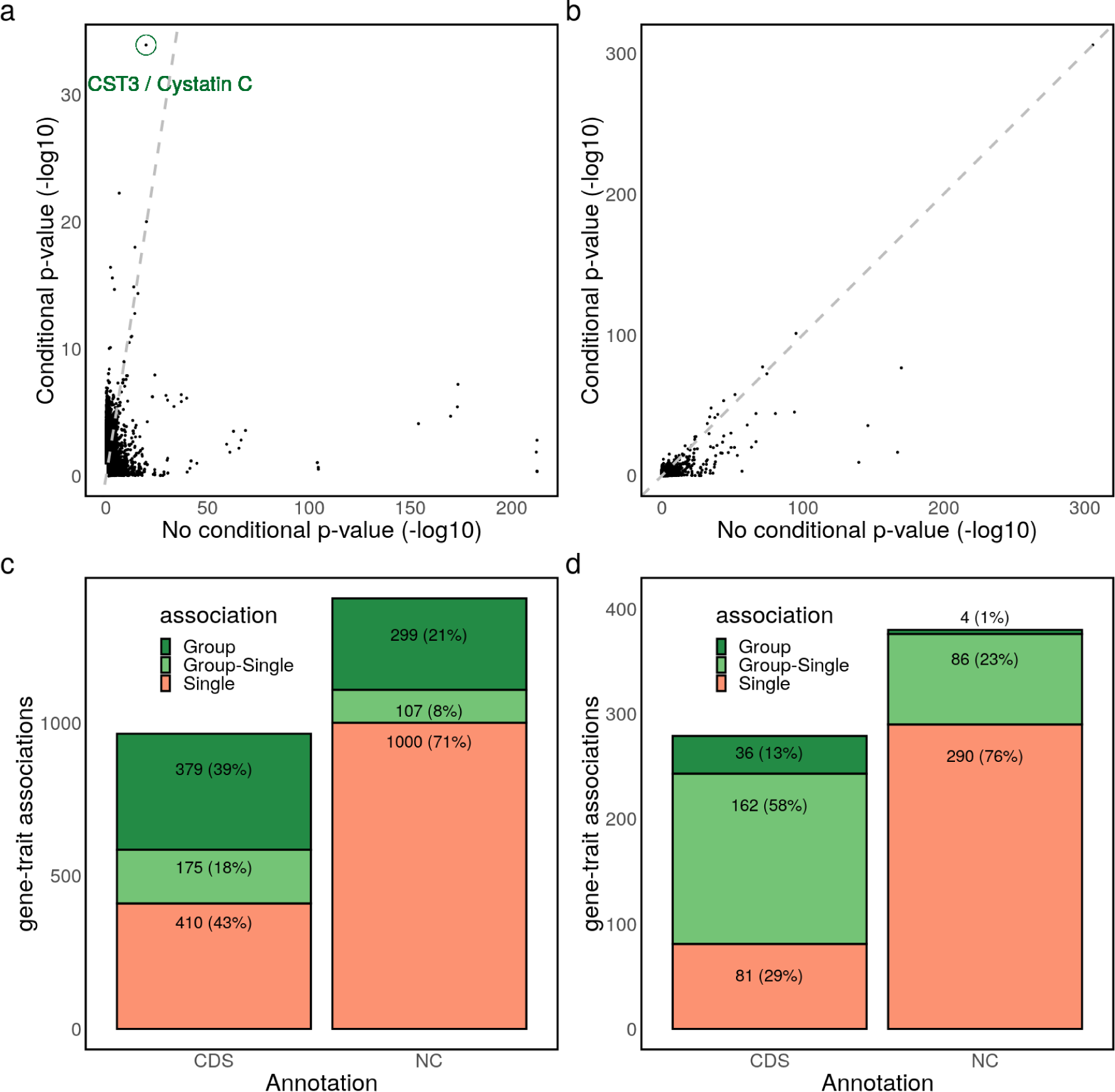
Rare variant association type and conditioning analysis. (a) NC annotation -log10 SKAT-O p-value comparison before and after conditioning on known common variants per trait; (b) same but for the CDS annotation. Note the difference in the y-axis range compared to the previous plot; (c) categorization of FDR<5% CDS and NC gene-trait associations into ‘single’, ‘group’ and ‘group-single’ associations based on comparison between single variant and collapsing analysis p-values; (d) categorization of gene-trait associations with P<1e^-9^.

For comparison, we performed conditional analysis on CDS gene-trait associations. We found between 38.5% and 39.8% associations retained at FDR<5% and P<1e^-9^, respectively (**Figure 4b**, **Figure 2g**). This constitutes a much higher proportion of retained associations compared to NC associations (9.5% and 3.7%). However, the decrease in significance of CDS associations when conditioning for common variants is substantially higher than reported in a previous UK Biobank rare variant exome study^10^ (see Discussion).

To further explore if LD is responsible for the drop in significance when conditioning for common variants, we focus on associations driven by a single variant, on which we can more easily study LD patterns. To first determine which associations found through collapsing approaches (SKAT-O) might be attributed to single variants, we performed single variant analysis for variants in NC and CDS annotations (see Methods). With this, we categorised significant gene-trait associations (FDR<5% or P<1e^-9^) from the collapsing analysis into three cases: (1) instances where no single variant annotated for the gene exhibited genome-wide significance (P<5e^-8^), indicating that only a collapsing test could unveil the association, (2) cases where at least one single variant association attained genome-wide significance, but the p-value was less significant than the SKAT-O p-value, suggesting that multiple variants may collectively contribute to the association, (3) situations in which at least one single variant association had a p-value equal to or stronger than the SKAT-O p-value (see Methods). Our findings indicated that a substantial proportion of NC gene-trait associations, ranging from 71% (FDR<5%) to 76% (P<1e^-9^), could potentially be attributed to a single variant alone (case 3, **Figure 4c,d**). Meanwhile, 8% (FDR<5%) to 23% (P<1e^-9^) of the gene-trait associations might be influenced by single variants or a combination of variants (case 2, **Supplementary Table 5**). About 21% of FDR<5% gene-trait associations were exclusively identifiable through collapsing approaches (case 1), but this proportion dropped to merely 1% for associations at P<1e^-9^. Notably, these patterns differed when considering CDS annotations, where a higher proportion of gene-trait associations could only be identified through collapsing approaches (39% at FDR<5%, 13% at P<1e^-9^). In fact, only 29% to 43% CDS associations appeared to be driven by a single variant alone (**Figure 4c,d**).

To understand the effect of LD between common and rare variants, we focused on a set of 245 single variants that were significant and sufficient in identifying NC gene-trait associations (FDR < 5%, **Figure 4c**). For each of the 245 single variants, we gathered GWAS conditional variants falling within a 1Mb window, yielding 163 single variants with a corresponding match and resulting in a total of 1006 “single-conditional” variant pairs. We then calculate LD between each pair of variants. Our analysis revealed that a staggering 99.4% of single variants (162 out of 163) were in LD with at least one conditional variant (D’ > 0.5, **Supplementary Table 6**, see Methods). We confirmed the effect of these conditional variants in LD, by performing conditional analysis for each single variant separately, conditioning only on those variants in LD. Accounting for conditional variants in LD was enough for 76.1% variant-traits associations to decrease significance (**Supplementary Figure 9, Supplementary Table 6**). Concretely, 49% of variant-trait associations with FDR < 5% no longer pass this threshold after conditioning, and 62.6% associations at P<1e^-9^ no longer pass this threshold. This demonstrates that LD between common and rare variants is a main driver of the observed associations and independent rare variant associations represent a minority of the associations.

Finally, we compared conditioning on genotypes of known variants, as done above, with using polygenic risk scores (PGS) as covariates. For this, we gathered publicly available PGS measurements for UK Biobank individuals across three traits (HDL cholesterol, LDL cholesterol and HbA1c)^46^. We found that accounting for PGS only slightly affects SKAT-O association p-values (**Supplementary Figure 10**, Spearman correlation = 0.92, p-value < 2.2e^-16^). In fact, PGS covariates actually increase association significance, with 93.1% gene-trait associations with P<1e^-9^ (without conditional) increasing in significance. In contrast, accounting for genotypes of independent GWAS hits (as above) for the same 3 traits only increases association significance in 10.3% of gene-trait associations with P<1e^-9^, with 89.7% of those associations no longer passing that significance threshold, consistent with our previous observations. These findings highlight a clear distinction between using PGS as covariates and conditioning on genotypes.

## Discussion

In this study, we leveraged biobank-scale WGS data and knowledge of gene regulation to tackle the challenge of linking non-coding rare variants to dozens of human traits. A few previous studies have proposed approaches to associate non-coding rare variants by aggregating variants in genomic sliding windows, functional annotations (e.g. enhancers)^47–49^ and/or variant deleteriousness regions^17,38^. However, these approaches have been applied to only a limited number of traits (usually fewer than 10) or used smaller sample sizes (<35 000 samples). Here, we combine variant deleteriousness filtering with multiple comprehensive datasets of gene regulation to investigate the association between non-coding rare variants and dozens of traits in >166 000 individuals. As consensus annotation of non-coding regions is lacking, we explore several orthogonal gene regulatory annotations in guiding variant region choice, thus decreasing the immense search-space inherent to the study of non-coding regions. Additionally, we aligned regulatory annotations with the specific cell types relevant to the traits under scrutiny, with the intuition that the use of biologically-relevant annotations is better suited to map causal loci. In this pilot study, we focused on studying blood-related phenotypes and regulatory annotations as these comprise some of the most extensively studied cell types and biomarkers, which are highly relevant across various diseases. Nevertheless, we consider our approach to be applicable to any cell type and human trait.

Our approach yielded a large number of non-coding gene-trait associations across the 59 traits, before conditioning for common variants (1406 gene-trait associations). We found this to be higher than for CDS regions (964 gene-trait associations) using the same models and approach. We note that we did not optimise discovery of CDS gene-trait associations, as we did not use coding variant annotations – such as missense and loss-of-function – which have been demonstrated to augment discovery of coding gene-trait associations^10,13^. Despite the higher count of gene-trait associations in non-coding regions, interpreting these associations presented challenges. Unlike coding regions, where a variant typically affects a single gene, regulatory variants (such as those in enhancer regions) might influence multiple nearby genes^25,41^. We accounted for this complexity in our methodology, allowing for several genes to be associated through the same variants. This approach inadvertently increased the number of associated genes and made it more challenging to distinguish causal genes from non-causal ones. Indeed, while we found that ∼28-54% of non-coding gene-trait associations were already documented GWAS catalog or Genebass databases, this proportion increased to nearly 100% when considering nearby genes within 1Mb. This suggests that our approach can unveil potentially causal regions and gene sets but often lack specificity in pinpointing individual causal genes. Although we found that nearly all our discovered gene-trait associations may already be documented, it is worth noting that the GWAS catalog and Genebass databases encompass results from the UK Biobank, based on SNP array and exome data, respectively. Our results thus suggest that employing WGS does not substantially enhance discovery yield, consistent with a recent study indicating little to no benefit in using WGS compared to a combination of SNP array, imputation, and exome sequencing within the UK Biobank^50^.

Along with the finding that the discovered non-coding gene-trait associations are known from GWAS or exome-seq rare variant studies, we found that only ∼9% of these are independent from known common variants. Indeed, we estimate that a large proportion of associated rare variants are in LD with GWAS common variants previously associated with the same trait. Previous studies on non-coding rare variants and lipid traits found that ∼32% signals are independent of common variants, although a more lenient association cutoff was used (P<1.2e^-3^, Bonferroni multiple test correction over 43 tests)^48^. A more recent approach on the same traits indicated that only ∼8% non-coding associations were independent, more in line with our results^51^. Regarding CDS regions, we observed that ∼40% CDS gene-trait associations are independent from known GWAS common variants, a large increase compared to non-coding regions. Analysis in lipid traits found between 66% and 75% coding rare variant associations being independent^48,51^. Moreover, previous studies on UK Biobank exome-seq conditioning on GWAS variants indicated that >90% of the CDS associations are independent^10^. Besides differences between non-coding and coding regions, the percentage of independent gene-trait associations may vary depending on the traits assessed, approach to conditioning (e.g. only nearby GWAS hits, or all hits in chromosome or genome), and the p-values cutoffs used to determine independence. For instance, we took a conservative approach by only considering a subset of 42 traits for which we had successfully obtained independent GWAS hits, whereas other studies might deem a rare variant association independent if no nearby GWAS hits are present^10^. Moreover, we performed conditional analysis across all possible gene-trait pairs and performed multiple test correction accordingly, whereas other studies only performed conditional analysis and multiple test correction on a subset of previously significant gene-trait associations^47,48,51^. Regarding traits, a study on 9 red blood phenotypes, which encompassed both coding and non-coding variants, determined that only 13% single variant-trait associations are independent from common variants^49^, suggesting estimates of independence may be lower for some blood cell traits. Furthermore, we found that accounting for common variants often decreases the signal from rare variants in LD, whereas the opposite is not true, as the common variant association may still be significant after accounting for the rare variant. However, this distinction may be confounded by allele frequencies and difference in power between common and rare variant analyses. Further research is required to fully determine whether the signal comes from the common or rare variants in LD.

When attempting to use PGS as covariates, instead of genotypes of GWAS variants, we observed a slight increase in the significance of association p-values. This aligns with findings of enhanced power in association testing with PGS covariates, a trend observed for both common variants^52^ and coding rare variants^53^. Our results indicate that this enhancement may extend to non-coding rare variant associations.

Variant selection is an important step in association analysis. Here, we leaned on CADD scores to focus on potentially deleterious variants. This genomic score is widely used in both coding and non-coding rare variant analysis^17,38,54^ and here allows us to compare coding and non-coding association results within the same framework. However, exploring other scores such as ReMM, GERP and Eigen^55–57^ for non-coding regions and AlphaMissense^58^ for coding regions, constitute an open avenue of research, with deleteriousness or pathogenicity models continuing to be developed. While using 150 000 WGS samples is a large increase in sample size compared to previous studies, another likely development is the continued increase in WGS samples in biobanks. Particularly noteworthy is the release of 500 000 whole genomes by the UK Biobank. Here we estimated that discovery power with both coding and non-coding rare variant analysis will clearly benefit from an increase in sample size. Furthermore, advancements in association testing are anticipated to include the integration of phasing information. This entails the incorporation of models designed to differentiate scenarios where one or both copies of a gene or regulatory element are impacted by rare variants. Notably, phasing information has been made available for the UK Biobank’s WGS dataset and is ready to be leveraged^59^.

In summary, we describe an approach to identify gene-trait associations with non-coding variants. Our findings indicate that these associations, for the most part, mirror known associations and are largely attributed to linkage disequilibrium between common and rare variants. This work thus serves as a cautionary tale in interpreting rare variant association results, as these should always be followed by robust conditional analyses. Further research is required to fully determine the relevance of non-coding rare variants in identifying independent trait associations, such as analyses on other traits, and increased sample size.

## Methods

### Ethics statement

This study relied on analyses of genetic data from the UKB cohort, which was collected with informed consent obtained from all participants. Data for this study were obtained under the UKB application licence number 66995. All data used in this research are publicly available to registered researchers through the UKB data-access protocol.

### Non-coding rare variant annotation

We considered three orthogonal non-coding rare variant annotations in blood cell types described below.

#### HIC (promoter capture Hi-C data, PCHiC)

This dataset consists of PCHiC data from 17 human primary hematopoietic cell types from Javierre et al. 2016^29^. We considered only intrachromosomal regions within 1Mb from the promoter region of a gene. Only PCHi-C interactions with CHiCAGO scores > 5 in at least one cell type were used.

#### CRD (cis-regulatory domains)

We obtained cis-regulatory domains derived from correlation between histone modification ChIP-seq peaks (H3K4me1 and H3K27ac) across 94 to 317 individuals and four blood cell types from Avalos et al. 2022^37^ (neutrophils, monocytes and T cells) and Delaneau et al. 2019^30^ (lymphoblastoid cell lines, LCLs). RNA-seq on the same samples was used in the original studies to link cis-regulatory domains to genes. We combined CRD-gene links across the four cell types with bedtools *merge* -o distinct (v2.26) to remove redundant regions per gene.

#### ABC (activity-by-contact model)

We obtained gene-enhancer association predictions from Nasser et al. 2021^34^ (‘AllPredictions.AvgHiC.ABC0.015.minus150.ForABCPaperV3.txt.gz’). We only considered interactions from a set of 60 blood cell types. Moreover, as done by the data providers, we only analysed interactions with ABC score > 0.1.

We further combined these three regulatory annotations into a single ‘NC’ (non-coding) annotation, comprising any variant falling in either of the three annotations above. For all non-coding annotations, to ensure regulatory association signals are not driven by coding regions, we excluded any overlap with CDS regions (as defined by Gencode v42) using bedtools *subtract* (v2.26). We used these same CDS regions as a comparison for association testing. In all analyses, we only considered autosomal protein coding genes based on Gencode v42, and excluded genes in the MHC region (chr6:28,510,120-33,480,577). When required, we lifted coordinates from hg19 to GRCh38 using the UCSC liftOver tool^60^ and converted gene symbols to Ensembl IDs using gProfiler (v105)^61^.

### UK Biobank whole genome sequencing and phenotypes

We use the 200K whole-genome sequencing dataset release from the UK Biobank, processed as described in the SHAPEIT Phasing Protocol for 200k WGS (field 20279). Briefly, starting with GraphTyper pVCFs, we excluded variants with (i) AAscore < 0.8, (ii) Hardy-Weinberg equilibrium (HWE) test p-value < 1e^-30^ (calculated in unrelated Caucasian samples), (iii) genotype missingness > 10% and (iv) excluded variant without a “PASS” in the VCF file. Multi-allelic variants were converted to bi-allelic sites using the bcftools *norm* (with -m -any parameters, v1.15).

We only considered 166 740 individuals of white British Caucasian ancestry (data field 22006) who met the following criteria: (i) reported and genetic sex is consistent (field 31 and 22001), (ii) had fewer than 10 third-degree relatives within the dataset (field 22021), and (iii) are not outliers in terms of heterozygosity or missing data rates (field 22027). We considered related samples, as these relationships are accounted for in association testing (see below).

We collected phenotypic data for a total of 59 blood-related traits, including 29 measures of blood cell counts and percentages, as well as 30 blood biomarkers (mean = 152 088 samples, **Supplementary Table 1**). In cases where multiple measurements of the same phenotype were available for a single individual, the first instance was used. We also collected age, sex and genotype principal components for each individual.

### Rare variant association testing

#### Variant selection

Prior to association testing, we filtered for UK Biobank WGS variants falling in any regulatory annotation position or CDS region with bcftools (v1.10.2). To test only variants predicted to be highly deleterious, we retained only variants with CADD (v1.6) score > 15 (‘All possible SNVs of hg38’)^39^. Of note, only CADD scores for single nucleotide variants were used, and thus only this type of variant was assessed in this study. VCF files were converted to PGEN with plink2 (--make-pgen erase-phase) for association testing.

#### Gene-based association testing

We performed genome-wide association tests using REGENIE (v3.2.8)^40^ for 166 740 individuals of white British Caucasian ancestry with WGS data. For REGINIE step 1, we included 360 166 UK Biobank SNP array (field 22418) variants for the same individuals, considering only variants in autosomes with MAF > 1%, <10% genotype missingness, HWE p-value > 1e^-15^, and pruned for LD with plink2 (parameter --indep-pairwise 500 50 0.2). The leave-one-chromosome-out (LOCO) models obtained with step 1 of REGENIE are used for step 2, for each trait. In step 2, we used sex (field 31), age (field 21022) and 10 ancestry-informative PCs derived from SNP array data (field 22009) as covariates. REGENIE step 2 for gene-based tests was run with the following parameters: --minMAC 1, --bsize 400, --aaf-bins 0.01, --vcf-tests skato, --vc-maxAAF 0.01, --apply-rint (i.e. rank-inverse normalise phenotype). Of note, this version of REGENIE collapses ultra-rare variants (defined as MAC < 10) into a single unit. We ran REGENIE step 2 separately for each annotation. Note that output of SKAT-O tests does not include estimates of beta (effect size) and its standard error, and thus these are not reported in this study.

#### Replication

To assess replication of association in different UKB samples, we split the 166 740 samples into 30 000 samples chosen randomly and the remaining 136 740 samples. From the 30 000 samples, we excluded those with at least one relative in the remaining set of 136 740 samples, retaining 26 212 samples. We then performed association separately on the set of 26 212 samples and the remaining 136 740 samples. In addition to these subsets of samples, gene-trait associations were also computed for a random subset of 83 370 samples (i.e. half of the 166 740 samples) to study the relationship between sample size and discovery power.

#### Single variant association testing

We performed association tests per variant against all 59 traits using REGENIE, from the same sets of variants used in gene-based tests. We ran Step 2 with the same parameters (except --vcf-tests skato), covariates and step 1 models as for gene-based tests. We included all single variants with MAF < 1%. Next, we compared association p-values between gene-based and single variant tests, to determine which associations are driven by one or more combinations of variants. For this, for each significant gene-trait association from gene-based tests (NC and CDS annotations), we determined whether at least one single variant linked to the same gene had a p-value as significant or more significant than the gene-based test (SKAT-O p-value). These cases were labelled as ‘Single cases’. For the remaining gene-trait associations (i.e. cases where gene-based association has higher significance than any single variant linked to the gene), we further categorised whether there is at least one single variant with p-value < 5e^-8^ (genome-wide significance threshold commonly used in single variant analysis), labelling these cases as ‘Group-single cases’. Finally, we labelled gene-trait associations where no single variant reaches this significance threshold as ‘Group cases’.

### Overlap with known associations

We evaluated how many of our gene-trait associations predictions overlap known gene-trait associations from two sources: (i) GWAS catalog^62^ (All associations v1.0, downloaded in March 2023) and (ii) Genebass^13^ based on UK Biobank exome-sequencing data. We obtained Genebass associations through Hail in August 2022 (‘results.mt’), and considered SKAT-O p-values below FDR 5% per trait for the ‘pLoF’ and ‘Missense|LC’ categories. All 59 traits had a match in both databases, with a total of 50 094 gene-trait pairs gathered. A gene-trait pair was deemed replicated if the pair was observed in at least one of the databases. The ‘EXPEC’ results were computed by shuffling once the SKAT-O p-values from the NC annotation, prior to FDR calculation and thus drawing random sets of gene-trait pairs matching the same number of discoveries. We used this to measure the amount of gene-trait associations across traits expected to be reproduced by chance.

To study groups of genes in a loci, we measured the overlap between associated and known gene-traits also when a gene within 500kb of each TSS region flank matches. For this, for each gene-trait pair, we gather all genes within this flanking region. If any of these genes have a known association to the same trait, we counted as a match. If neither the current gene or nearby genes matches, a match is not counted.

### Association conditional analysis

To condition for known common variants associated with traits, we first obtained GWAS summary statistics for the EUR population from the Pan UK Biobank (Pan-UKB team. https://pan.ukbb.broadinstitute.org. 2020). These GWAS computations encompassed the samples used in our study, as well as others with SNP array information (mean = 369 748 samples for the traits assessed). We then obtained independently associated common variants for each trait by using the GCTA (v1.94.1) *cojo* joint model function^44^, with parameters: window = 10 Mb, collinearity = 0.9, for all GWAS variants with P < 1e^-7^. To run GCTA *cojo* models, we used LD structure estimates for GWAS common variants (WTCHG imputed SNP array data, field 2282) from 10 000 UK Biobank samples randomly sampled from the set of white British individuals with WGS data. We used variants with P < 1e^-7^ (instead of the typically used P < 5e^-8^ cutoff) to ensure that all known common variant signals are accounted for. Finally, performed conditional analysis for genotypes of independent GWAS hits (SNP array data, field 2282) using REGENIE ---condition-list and --condition-file. Unless stated otherwise, we used all independent GWAS hits across all chromosomes associated with a trait together when conditioning. We kept all other parameters and REGENIE step1 models as in the analysis without conditioning. We performed conditional analyses in a subset of 42 traits out of the 59 tested, due to the inability of finding all independent common variants for the remaining traits. We only show results of conditional analysis for those 42 traits for which we performed a full conditional analysis. We re-estimated the FDR (Benjamini-Hochberg) of SKAT-O p-values for the set of 42 traits.

We calculated linkage disequilibrium (LD) R2 and D’ metrics using the plink2 --ld flag. First, we lifted SNP array hg19 coordinates to hg38 assembly with the UCSC liftOver tool. Then, for each of 245 rare variants associated with a trait at FDR<5%, we gathered all independent GWAS common variants (from GCTA *cojo*) falling within a 1Mb window. A total of 163 unique rare variants had at least one common variant within this window. We measured LD between 1006 rare and common variants paired in this manner.

In addition, we obtained polygenic risk scores (PGS) for three traits (LDL, HDL and HbA1c) from Thompson et al. 2022^46^ available in the UK Biobank RAP (standard PGS, fields 26238, 26242 and 26250). These PGS are based on GWAS external to the UK Biobank. We used the PGS for each trait as a covariate when performing REGENIE SKAT-O association tests, for each trait separately.

## Data availability

The lists of gene-trait associations identified here are available as Supplementary Tables. The phased WGS data and individual phenotypes can be accessed via the UKB Research Analysis Platform (RAP): https://ukbiobank.dnanexus.com/landing. This platform is open to researchers who are listed as collaborators on UKB-approved access applications.

## Code availability

Code and source data to reproduce analysis and plots have been deposited in a github repository: https://github.com/diogomribeiro/noncoding_rvat.

## Acknowledgments

This work was funded by a Swiss National Science Foundation (SNSF) project grant (PP00P3_176977). The funders had no role in study design, data collection and analysis, decision to publish or preparation of the manuscript. We thank Zoltán Kutalik and Robin Hofmeister for fruitful discussions.

## Author contributions

D.M.R. performed the experiments and analysed the data. D.M.R. wrote the manuscript with inputs from O.D. D.M.R. and O.D. conceived and designed the study. O.D. supervised the study.

## Supplementary Information

**Supplementary Table 3.**
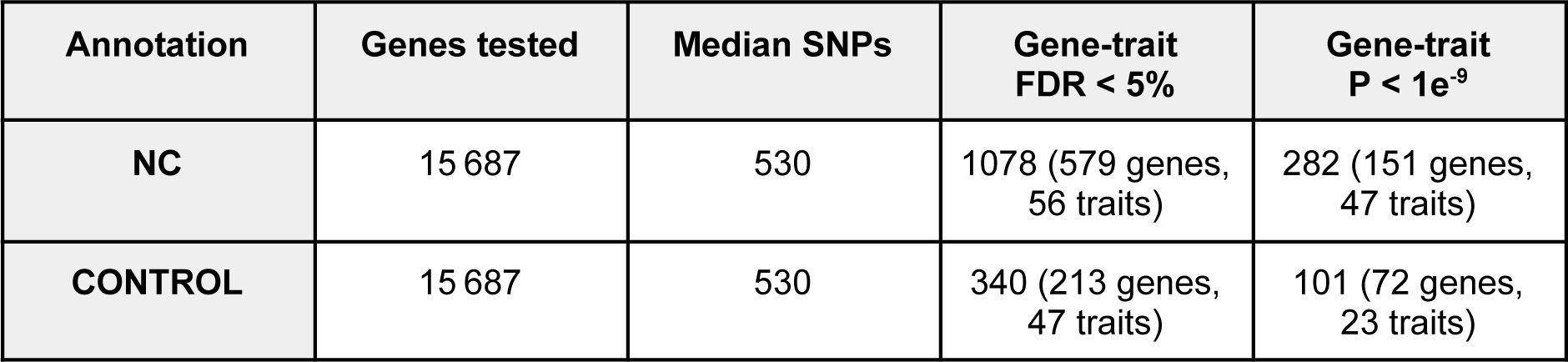
Summary of association tests between NC and CONTROL annotations. Only genes tested in both annotations were assessed. The number of SNPs per gene are matched between the annotations.

**Supplementary Figure 1.**
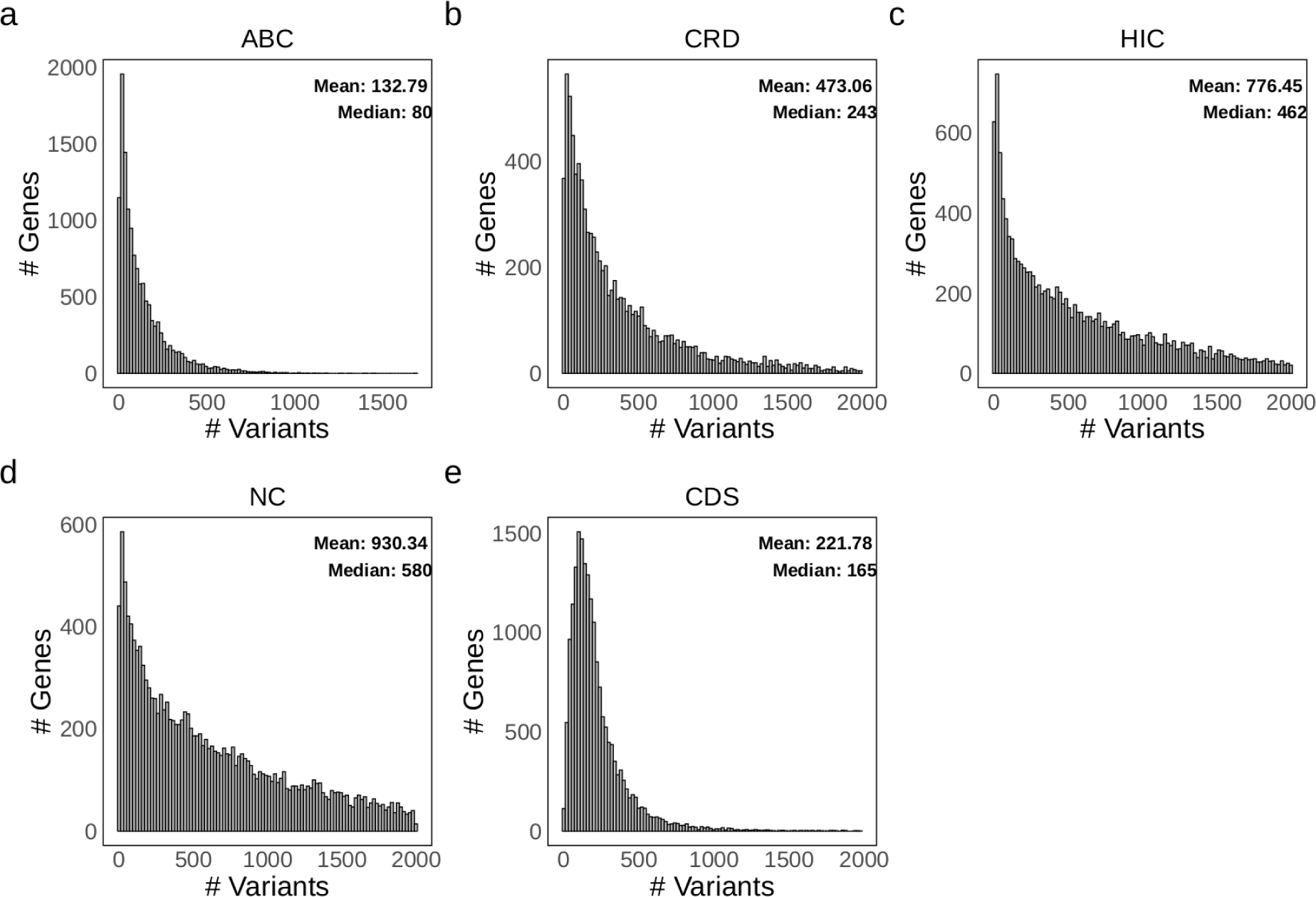
Number of rare variants per annotation. This refers to the number of variants after CADD > 15 filter and exclusion of CDS regions (except for CDS annotation). NC is the non-redundant combination of variants from ABC, CRD and HIC. X-axis is capped at 2000 for visibility, but the median and mean were calculated before capping.

**Supplementary Figure 2.**
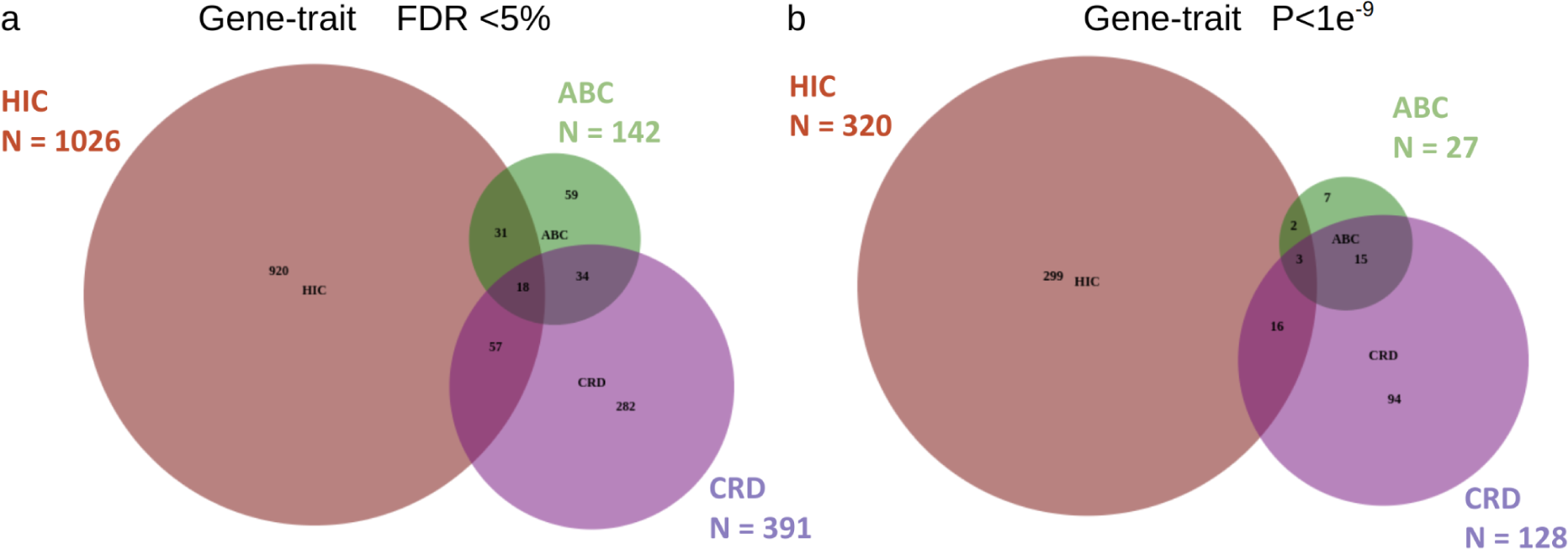
Venn diagram of gene-trait associations between non-coding annotations. (a) associations with FDR<5% and (b) associations with P<1e^-9^. Produced with DeepVenn.com (2020 Tim Hulsen).

**Supplementary Figure 3.**
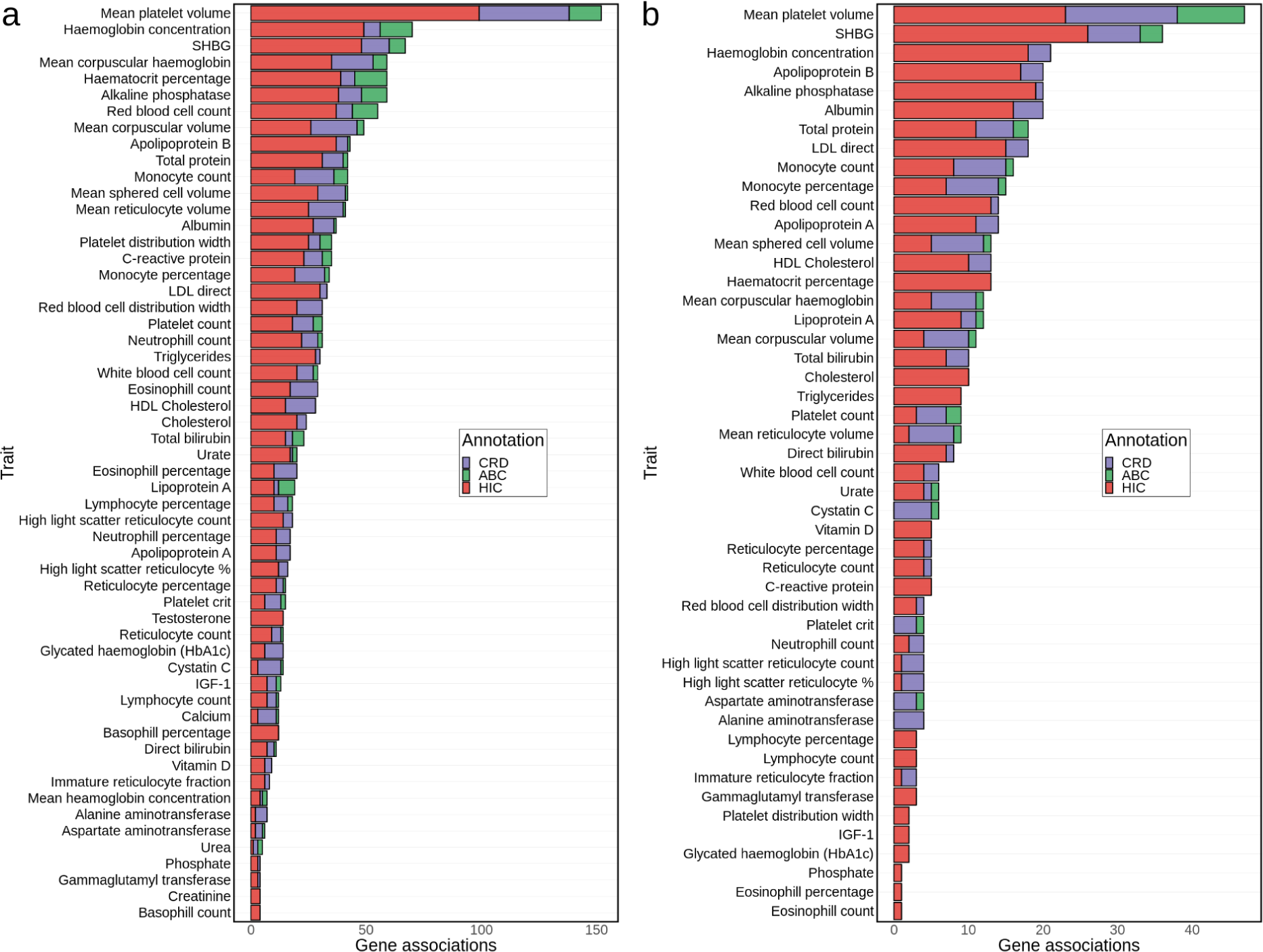
Number of genes associated per trait depending on the non-coding annotation. (a) SKAT-O associations with FDR < 5%, (b) SKAT-O associations with P<1e^-9^. Note that the same gene may be associated through more than one annotation, and thus is counted multiple times.

**Supplementary Figure 4.**
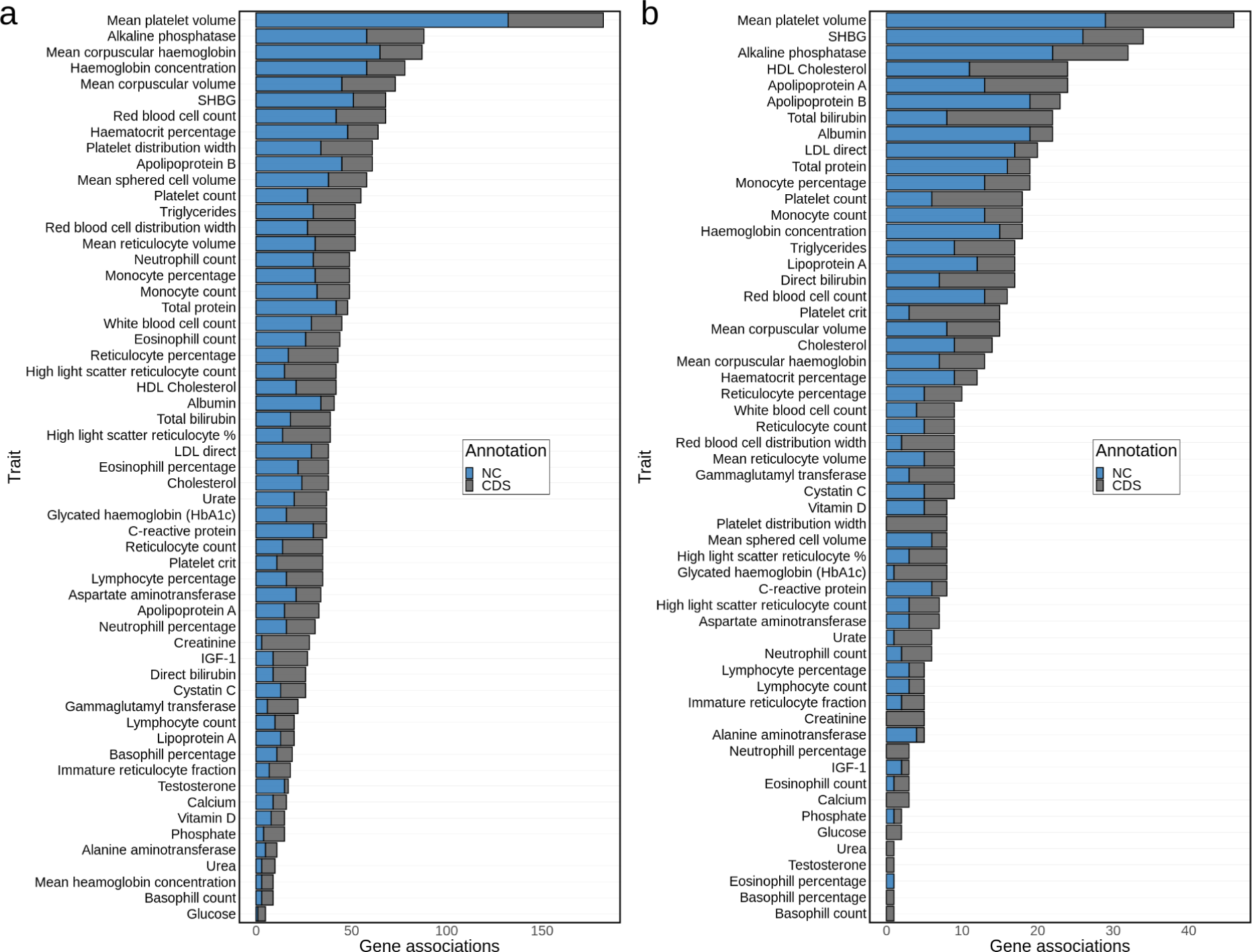
Number of genes associated per trait between NC and CDS annotations. (a) SKAT-O associations with FDR < 5%, (b) SKAT-O associations with P<1e^-9^. Note that the same gene may be associated through more than one annotation, and thus is counted multiple times.

**Supplementary Figure 5.**
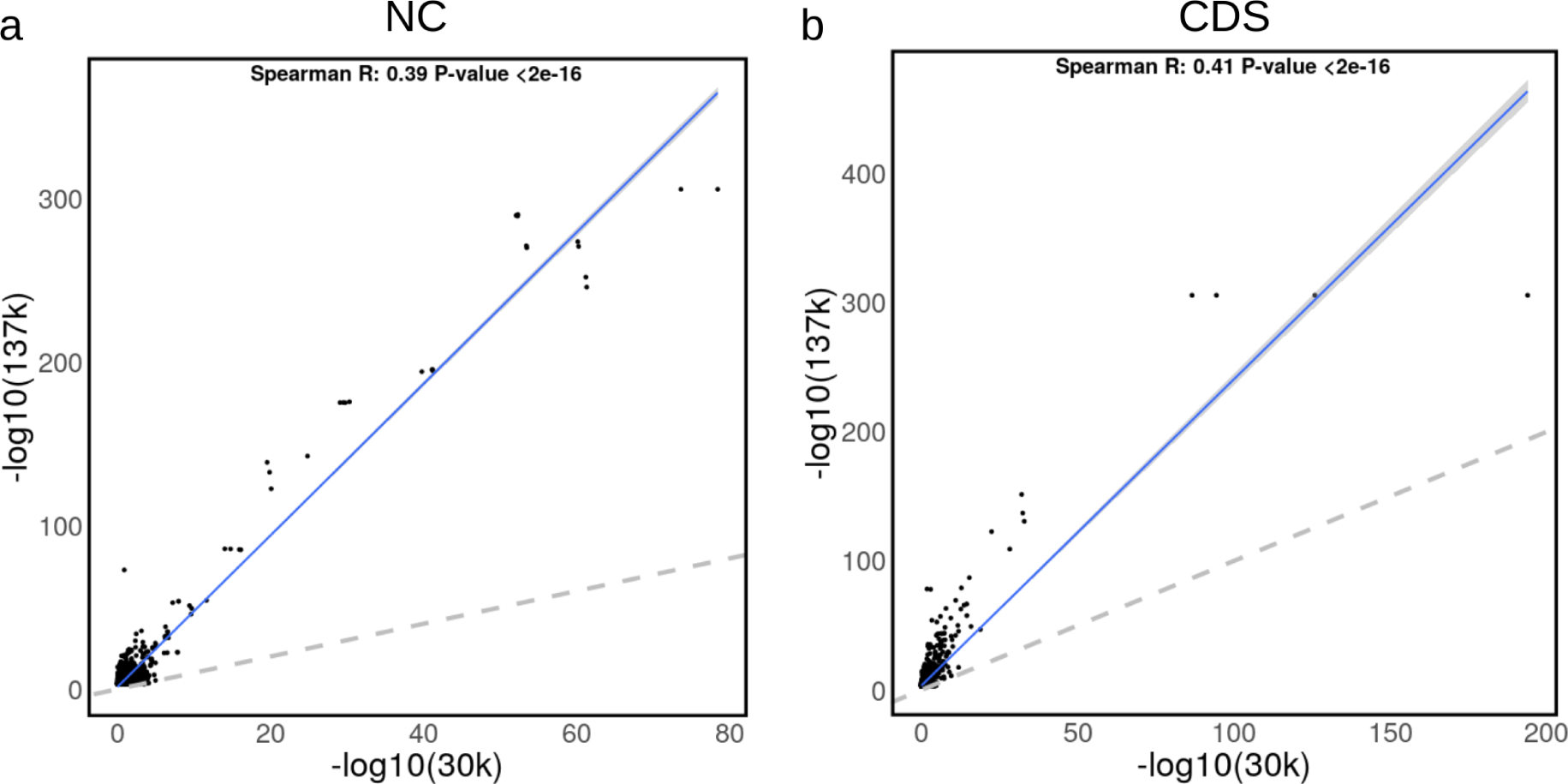
Correlation of association p-values between two subsets of samples (137k samples and 30k samples) (a) -log_10_ SKAT-O p-values for NC annotation associations; (b) -log_10_ SKAT-O p-values for CDS annotation associations. Associations with -log_10_ p-value < 3 in the 137k samples are not displayed.

**Supplementary Figure 6.**
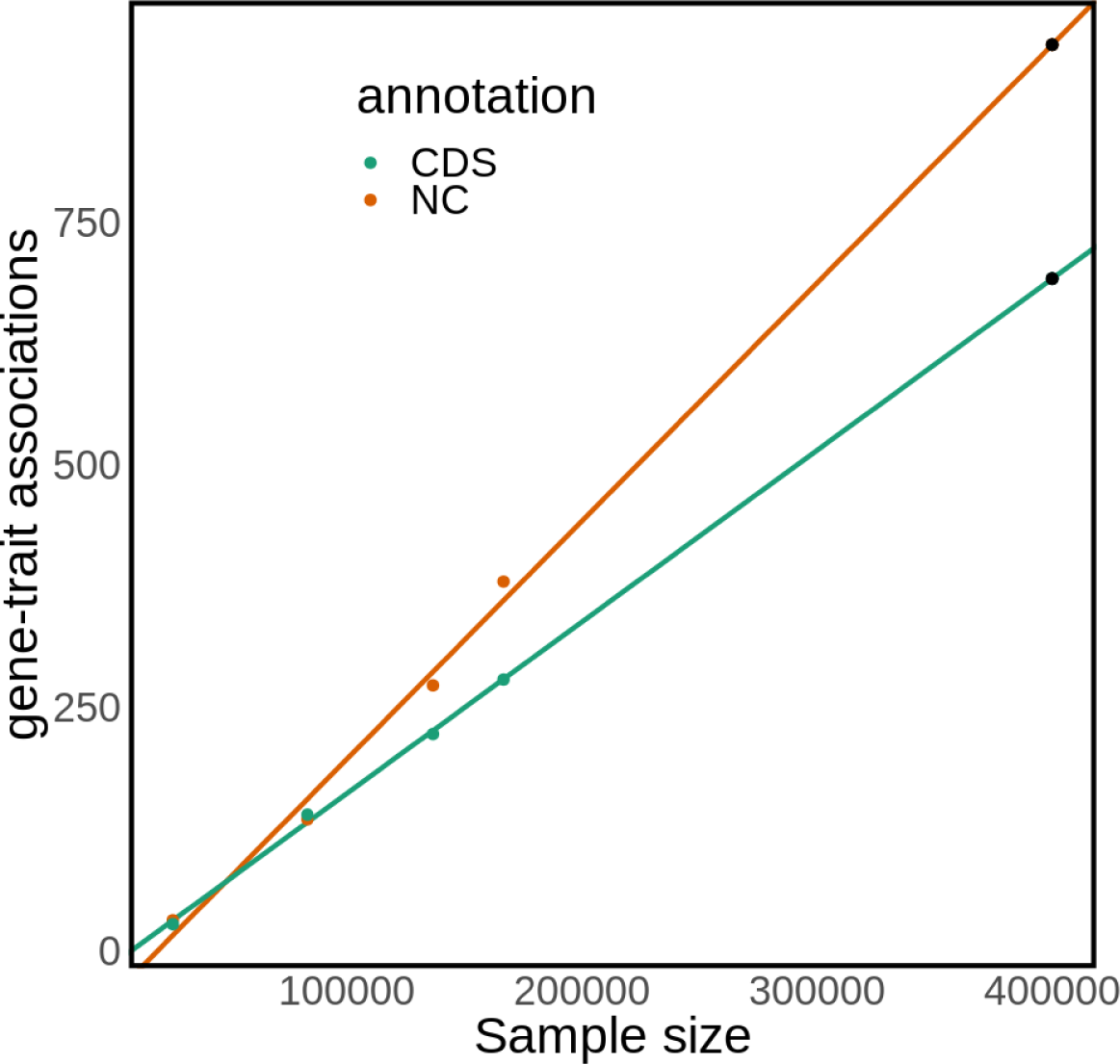
Correspondence between sample size and number of discovered gene-trait associations at P<1e^-9^. The black dot at 400 000 samples is a prediction based on the previous four samples sizes.

**Supplementary Figure 7.**
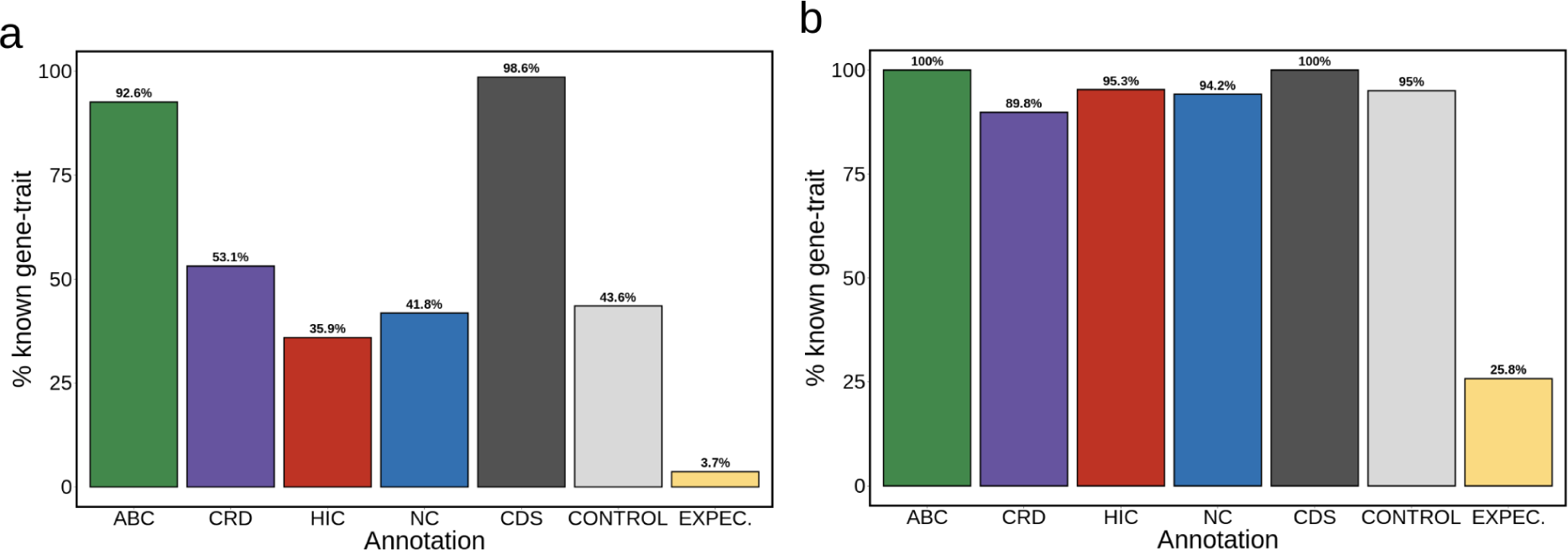
Percentage of gene-trait pairs with P<1e^-9^ matching known gene-trait associations. (a) direct gene-trait match; (b) direct gene-trait associations or indirect associations within a 1Mb window. Known associations considered if present either in GWAS catalog or Genebass datasets. ‘EXPEC.’ refers to random shuffling of NC annotation association p-values.

**Supplementary Figure 8.**
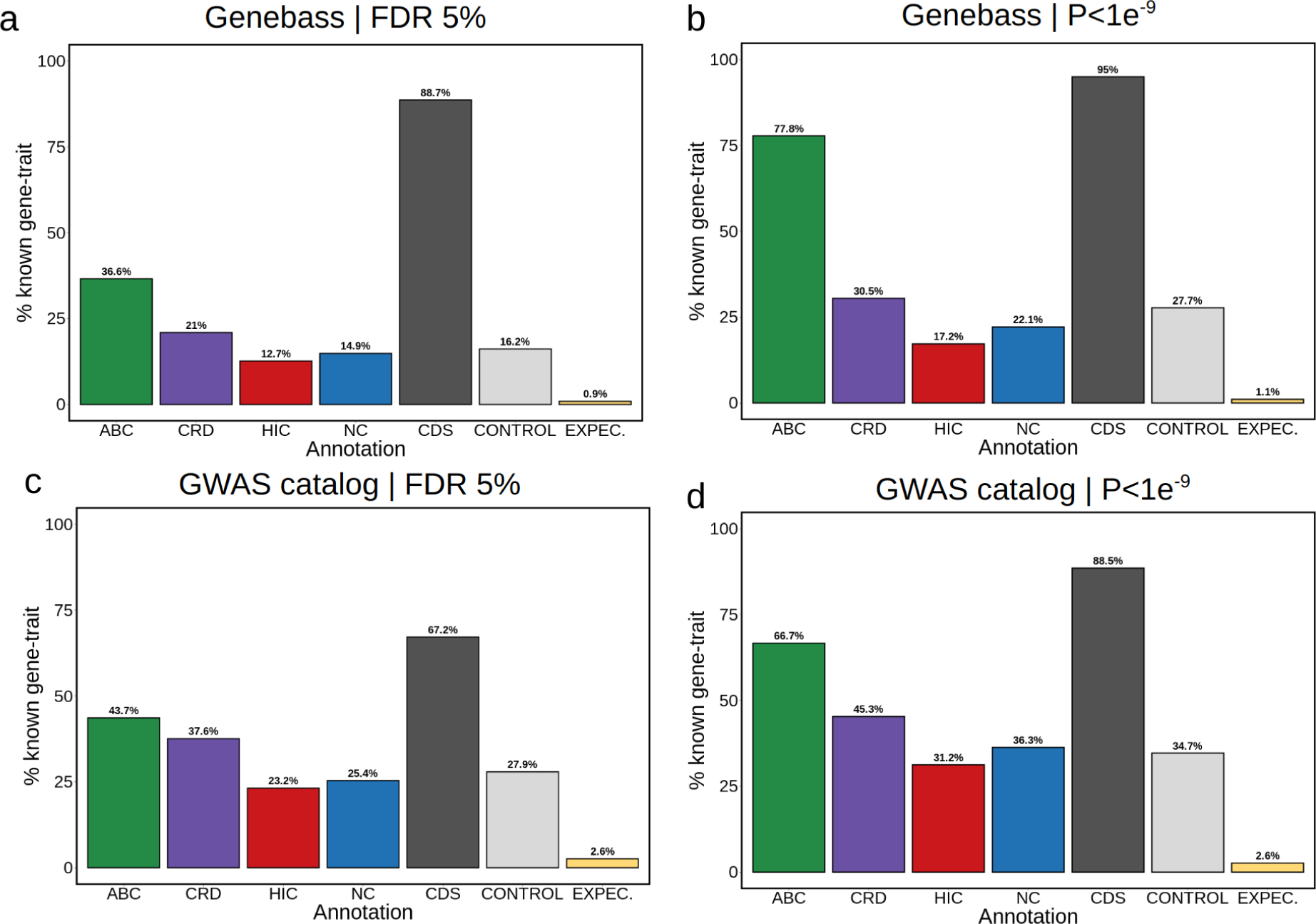
Percentage of gene-trait pairs matching known gene-trait associations depending on database. (a) discovery at FDR<5% and replication in Genebass database; (b) discovery at P<1e^-9^ significance and Genebass database; (c) discovery at FDR<5% and replication in the GWAS catalog; (d) discovery at P<1e^-9^ and replication in the GWAS catalog. ‘EXPEC.’ refers to random shuffling of NC annotation association p-values.

**Supplementary Figure 9.**
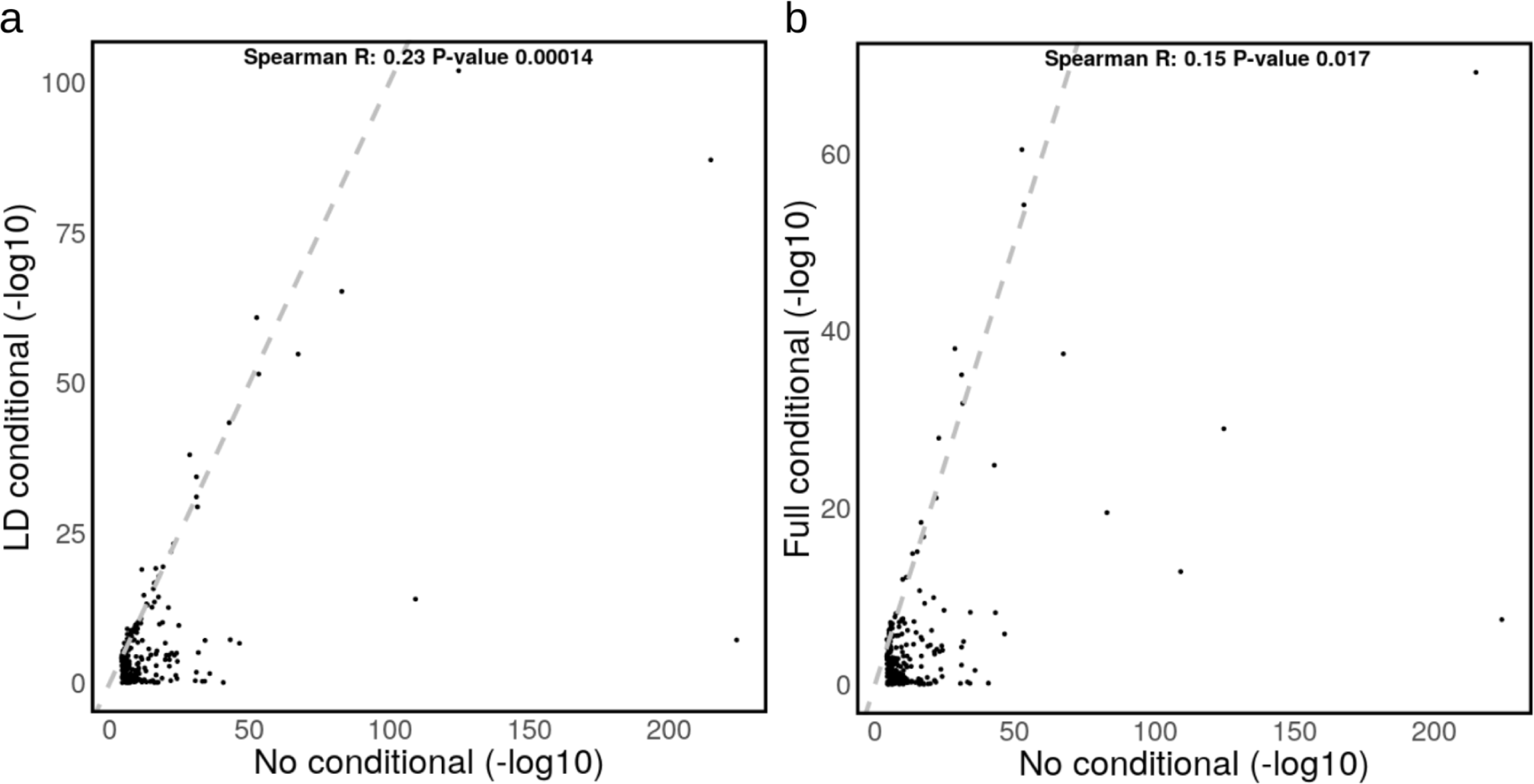
Association p-values conditioning for sets of common variants. 259 single rare variant associations with traits were tested for association under different conditioning models. (a) comparison of association p-values between no conditioning (x-axis) and conditioning for GWAS common variants in a 1Mb window that are in LD (D’>0.5) with the rare variant (y-axis); (b) comparison of association p-values between no conditioning (x-axis) and conditioning for all GWAS common variants (y-axis). Cases where -log_10_ p-value < 1 are not displayed, but are included for correlation.

**Supplementary Figure 10.**
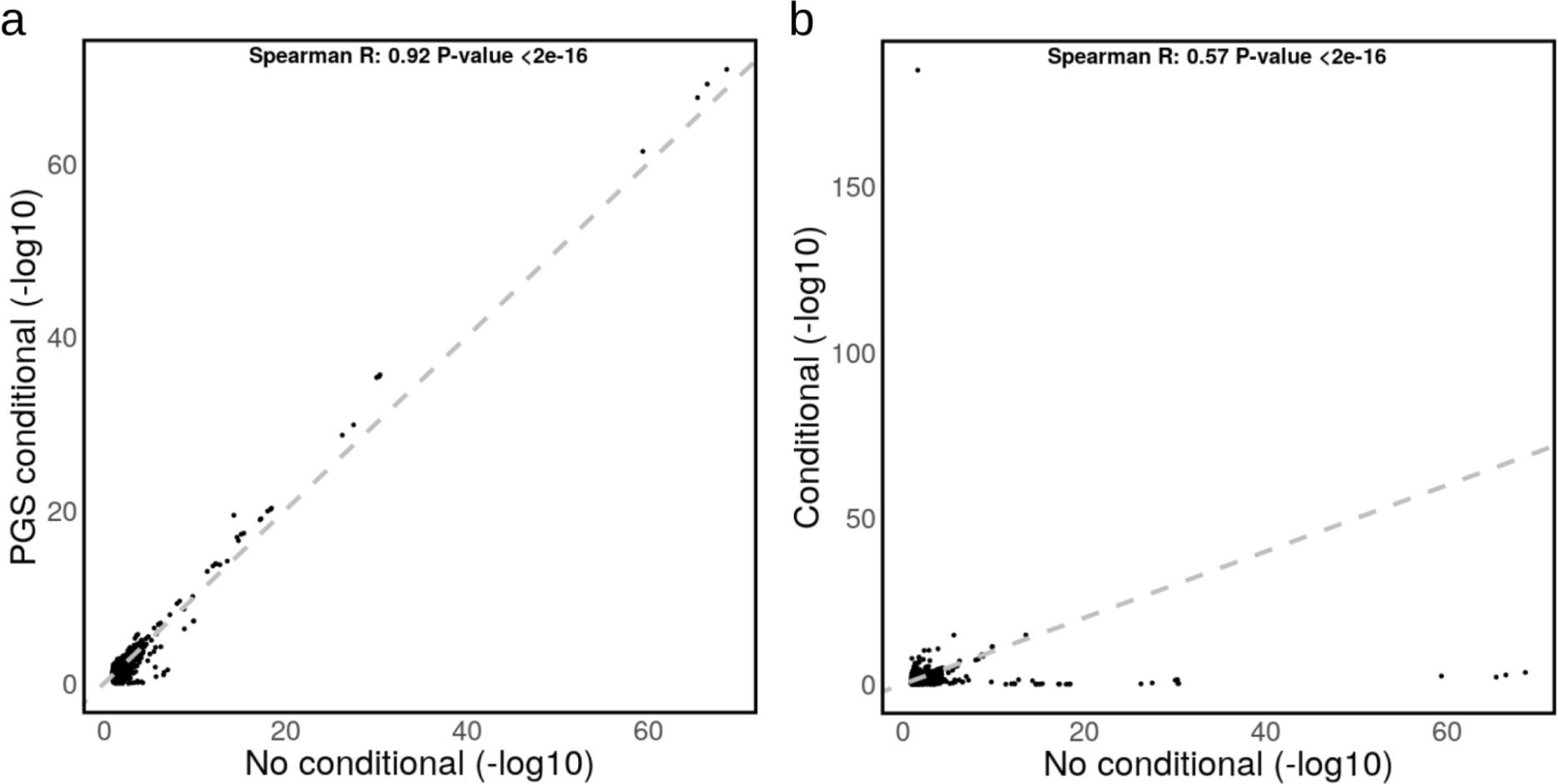
SKAT-O association p-values in three traits with PGS as covariates. (a) comparison of NC association p-values between no conditioning (x-axis) and using PGS for each trait independently as covariate (y-axis); (b) comparison of NC association p-values between no conditioning (x-axis) and conditioning for independent GWAS hits (y-axis). Cases where -log_10_ p-value < 1 are not displayed, but are included for correlation.

## Bibliography

1. Abdellaoui, A., Yengo, L., Verweij, K. J. H. & Visscher, P. M. 15 years of GWAS discovery: Realizing the promise. Am. J. Hum. Genet. 110, 179–194 (2023).

2. Rubinacci, S., Delaneau, O. & Marchini, J. Genotype imputation using the Positional Burrows Wheeler Transform. PLoS Genet. 16, e1009049 (2020).

3. Li, Y., Willer, C., Sanna, S. & Abecasis, G. Genotype imputation. Annu. Rev. Genomics Hum. Genet. 10, 387–406 (2009).

4. Zuk, O., Hechter, E., Sunyaev, S. R. & Lander, E. S. The mystery of missing heritability: Genetic interactions create phantom heritability. Proc. Natl. Acad. Sci. U. S. A. 109, 1193–1198 (2012).

5. Wainschtein, P. et al. Assessing the contribution of rare variants to complex trait heritability from whole-genome sequence data. Nat. Genet. 54, 263–273 (2022).

6. Zuk, O. et al. Searching for missing heritability: designing rare variant association studies. Proc. Natl. Acad. Sci. U. S. A. 111, E455–64 (2014).

7. Claussnitzer, M. et al. A brief history of human disease genetics. Nature 577, 179–189 (2020).

8. Wang, Q. et al. Rare variant contribution to human disease in 281,104 UK Biobank exomes. Nature 597, 527–532 (2021).

9. Momozawa, Y. & Mizukami, K. Unique roles of rare variants in the genetics of complex diseases in humans. J. Hum. Genet. 66, 11–23 (2021).

10. Backman, J. D. et al. Exome sequencing and analysis of 454,787 UK Biobank participants. Nature 599, 628–634 (2021).

11. Hall, S. S. Genetics: a gene of rare effect. Nature 496, 152–155 (2013).

12. Sabatine, M. S. et al. Evolocumab and Clinical Outcomes in Patients with Cardiovascular Disease. N. Engl. J. Med. 376, 1713–1722 (2017).

13. Karczewski, K. J. et al. Systematic single-variant and gene-based association testing of thousands of phenotypes in 394,841 UK Biobank exomes. Cell Genom 2, 100168 (2022).

14. Hindorff, L. A. et al. Potential etiologic and functional implications of genome-wide association loci for human diseases and traits. Proc. Natl. Acad. Sci. U. S. A. 106, 9362–9367 (2009).

15. Maurano, M. T. et al. Systematic localization of common disease-associated variation in regulatory DNA. Science 337, 1190–1195 (2012).

16. Hawkes, G., et al. Whole genome sequencing analysis identifies rare, large-effect non-coding variants and regions associated with circulating protein levels. bioRxiv (2023) doi:10.1101/2023.11.04.565589.

17. Hawkes, G., et al. Whole genome association testing in 333,100 individuals across three biobanks identifies rare non-coding single variant and genomic aggregate associations with height. bioRxiv (2023) doi:10.1101/2023.11.19.566520.

18. Bocher, O. & Génin, E. Rare variant association testing in the non-coding genome. Hum. Genet. 139, 1345–1362 (2020).

19. Bycroft, C. et al. The UK Biobank resource with deep phenotyping and genomic data. Nature 562, 203–209 (2018).

20. Halldorsson, B. V. et al. The sequences of 150,119 genomes in the UK biobank. Nature 607, 732–740 (2022).

21. Taliun, D. et al. Sequencing of 53,831 diverse genomes from the NHLBI TOPMed Program. Nature 590, 290–299 (2021).

22. All of Us Research Program Investigators, et al. The ‘All of Us’ Research Program. N. Engl. J. Med. 381, 668–676 (2019).

23. Andersson, R. & Sandelin, A. Determinants of enhancer and promoter activities of regulatory elements. Nat. Rev. Genet. 21, 71–87 (2020).

24. Claringbould, A. & Zaugg, J. B. Enhancers in disease: molecular basis and emerging treatment strategies. Trends Mol. Med. 27, 1060–1073 (2021).

25. Ribeiro, D. M. et al. The molecular basis, genetic control and pleiotropic effects of local gene co-expression. Nat. Commun. 12, 4842 (2021).

26. Hoellinger, T. et al. Enhancer/gene relationships: need for more reliable genome-wide reference sets. Front. Bioinform. 3, 1092853 (2023).

27. Sonawane, A. R. et al. Understanding Tissue-Specific Gene Regulation. Cell Rep. 21, 1077–1088 (2017).

28. Lieberman-Aiden, E. et al. Comprehensive mapping of long-range interactions reveals folding principles of the human genome. Science 326, 289–293 (2009).

29. Javierre, B. M. et al. Lineage-Specific Genome Architecture Links Enhancers and Non-coding Disease Variants to Target Gene Promoters. Cell 167, 1369–1384.e19 (2016).

30. Delaneau, O. et al. Chromatin three-dimensional interactions mediate genetic effects on gene expression. Science 364, (2019).

31. Roadmap Epigenomics Consortium et al. Integrative analysis of 111 reference human epigenomes. Nature 518, 317–330 (2015).

32. Boix, C. A., James, B. T., Park, Y. P., Meuleman, W. & Kellis, M. Regulatory genomic circuitry of human disease loci by integrative epigenomics. Nature 590, 300–307 (2021).

33. Fulco, C. P. et al. Activity-by-contact model of enhancer-promoter regulation from thousands of CRISPR perturbations. Nat. Genet. 51, 1664–1669 (2019).

34. Nasser, J. et al. Genome-wide enhancer maps link risk variants to disease genes. Nature 593, 238–243 (2021).

35. Gazal, S. et al. Combining SNP-to-gene linking strategies to identify disease genes and assess disease omnigenicity. Nat. Genet. 54, 827–836 (2022).

36. Dey, K. K. et al. SNP-to-gene linking strategies reveal contributions of enhancer-related and candidate master-regulator genes to autoimmune disease. Cell Genom 2, (2022).

37. Avalos, D. et al. Genetic variation in cis-regulatory domains suggests cell type-specific regulatory mechanisms in immunity. Commun Biol 6, 335 (2023).

38. Bocher, O. et al. Testing for association with rare variants in the coding and non-coding genome: RAVA-FIRST, a new approach based on CADD deleteriousness score. PLoS Genet. 18, e1009923 (2022).

39. Rentzsch, P., Witten, D., Cooper, G. M., Shendure, J. & Kircher, M. CADD: predicting the deleteriousness of variants throughout the human genome. Nucleic Acids Res. 47, D886–D894 (2019).

40. Mbatchou, J. et al. Computationally efficient whole-genome regression for quantitative and binary traits. Nat. Genet. 53, 1097–1103 (2021).

41. M Ribeiro, D., Ziyani, C. & Delaneau, O. Shared regulation and functional relevance of local gene co-expression revealed by single cell analysis. Commun Biol 5, 876 (2022).

42. Hambleton, S. et al. IRF8 mutations and human dendritic-cell immunodeficiency. N. Engl. J. Med. 365, 127–138 (2011).

43. Tamura, T., Kurotaki, D. & Koizumi, S.-I. Regulation of myelopoiesis by the transcription factor IRF8. Int. J. Hematol. 101, 342–351 (2015).

44. Yang, J. et al. Conditional and joint multiple-SNP analysis of GWAS summary statistics identifies additional variants influencing complex traits. Nat. Genet. 44, 369–75, S1–3 (2012).

45. Vuckovic, D. et al. The Polygenic and Monogenic Basis of Blood Traits and Diseases. Cell 182, 1214–1231.e11 (2020).

46. Thompson, D. J., et al. UK Biobank release and systematic evaluation of optimised polygenic risk scores for 53 diseases and quantitative traits. bioRxiv (2022) doi:10.1101/2022.06.16.22276246.

47. Li, Z. et al. A framework for detecting noncoding rare-variant associations of large-scale whole-genome sequencing studies. Nat. Methods 19, 1599–1611 (2022).

48. Li, X. et al. Powerful, scalable and resource-efficient meta-analysis of rare variant associations in large whole genome sequencing studies. Nat. Genet. 55, 154–164 (2023).

49. Hu, Y. et al. Whole-genome sequencing association analysis of quantitative red blood cell phenotypes: The NHLBI TOPMed program. Am. J. Hum. Genet. 108, 874–893 (2021).

50. Gaynor, S. M., et al. Yield of genetic association signals from genomes, exomes, and imputation in the UK biobank. bioRxiv (2023) doi:10.1101/2023.09.13.23295479.

51. Li, X., et al. A statistical framework for powerful multi-trait rare variant analysis in large-scale whole-genome sequencing studies. bioRxiv (2023) doi:10.1101/2023.10.30.564764.

52. Bennett, D., O’Shea, D., Ferguson, J., Morris, D. & Seoighe, C. Controlling for background genetic effects using polygenic scores improves the power of genome-wide association studies. Sci. Rep. 11, 19571 (2021).

53. Jurgens, S. J. et al. Adjusting for common variant polygenic scores improves yield in rare variant association analyses. Nat. Genet. 55, 544–548 (2023).

54. Dutta, D. et al. Meta-MultiSKAT: Multiple phenotype meta-analysis for region-based association test. Genet. Epidemiol. 43, 800–814 (2019).

55. Smedley, D. et al. A Whole-Genome Analysis Framework for Effective Identification of Pathogenic Regulatory Variants in Mendelian Disease. Am. J. Hum. Genet. 99, 595–606 (2016).

56. Davydov, E. V. et al. Identifying a high fraction of the human genome to be under selective constraint using GERP++. PLoS Comput. Biol. 6, e1001025 (2010).

57. Ionita-Laza, I., McCallum, K., Xu, B. & Buxbaum, J. D. A spectral approach integrating functional genomic annotations for coding and noncoding variants. Nat. Genet. 48, 214–220 (2016).

58. Cheng, J. et al. Accurate proteome-wide missense variant effect prediction with AlphaMissense. Science 381, eadg7492 (2023).

59. Hofmeister, R. J., Ribeiro, D. M., Rubinacci, S. & Delaneau, O. Accurate rare variant phasing of whole-genome and whole-exome sequencing data in the UK Biobank. Nat. Genet. 55, 1243–1249 (2023).

60. Lee, B. T. et al. The UCSC Genome Browser database: 2022 update. Nucleic Acids Res. 50, D1115–D1122 (2022).

61. Raudvere, U. et al. g:Profiler: a web server for functional enrichment analysis and conversions of gene lists (2019 update). Nucleic Acids Res. 47, W191–W198 (2019).

62. Sollis, E. et al. The NHGRI-EBI GWAS Catalog: knowledgebase and deposition resource. Nucleic Acids Res. 51, D977–D985 (2023).

